# Extracellular vesicles aid in the transfer of long-term associative memory between *Caenorhabditis elegans*

**DOI:** 10.1101/2025.02.26.640282

**Authors:** Monmita Bhar, Tanumoy Nandi, Hari Narayanan, Kamal Kishore, Kavita Babu

## Abstract

Memory formation is necessary for the survival of animals across phyla. Here, we elucidate the mechanism underlying the formation of long-term associative memory (LTAM) formed by treating *Caenorhabditis elegans* with a volatile chemoattractant and heat. Previous work has shown that training animals with a paradigm involving heat and isoamyl alcohol (IAA) simultaneously, causes *C. elegans* to lose their attraction to IAA. In this study, we elaborate on the mechanism behind this LTAM formation and suggest that during training with heat and IAA, *C. elegans* release extracellular vesicles (EVs) that upon being taken up by the same trained animals or their untrained counterparts causes the organism to lose attraction to IAA. Our data suggests that the vesicles are highly specific to the training paradigms used and differ with differing cues. Finally, we show that this mechanism of transfer of LTAM appears to be conserved between *C. elegans* and *C. briggsae* allowing for both intra and interspecies transfer of memory.

## Introduction

Native preferences and innate behaviours are essential for the survival of living organisms. Many innate behaviours represent the hardwired neural circuitry put in place during the development of an animal (Bateson & Mameli, 2007; Tierney, 1986). *Caenorhabditis elegans* has been widely used to study behavioural plasticity because of its ability to learn and modify its innate behaviour (reviewed in (Ardiel & Rankin, 2010)). With a nervous system consisting of 302 neurons, these animals show a variety of robust and adaptable behaviours like chemotaxis and thermotaxis (reviewed in (Ardiel & Rankin, 2010; Hart, 2006)). These behaviours allow *C. elegans* to move towards food sources, avoid toxic bacteria, and find mates, all of which rely heavily on chemosensation by the animal and innate preferences for certain odours ((Ballestriero *et al*, 2016) and reviewed in (Ferkey *et al*, 2021)). For example, under well fed naïve condition, *C. elegans* are attracted to odorants like isoamyl alcohol (IAA), which can be sensed by the AWC neuron (reviewed in (Bargmann, 2006)). However, this odour to behaviour relationship can be modified if the animal is exposed to periods of starvation along with IAA (Pereira & van der Kooy, 2012). Similarly, neutral concentrations of butanone can be turned attractive when repetitively paired with food. These modified behaviours can persist for up to forty hours post training (Kauffman *et al*, 2011). The above examples indicate the formation of associative memory, that is brought about as a result of the animal’s interaction with two simultaneous cues.

Associative memory in *C. elegans* can be classified as short-term or long-term associative memory, depending on how long it can last. Previous studies have shown that pairing cues for a single conditioning period (mass training) forms short-term associative memory (STAM) in *C. elegans*, that last for around 2 hours. However, multiple conditioning along with periods of rest in between forms long-term associative memory (LTAM), which can last from 20 to 40 hours (Kauffman *et al*., 2011). LTAM formation in *C. elegans* has largely been examined by pairing the presence/absence of food (unconditioned stimulus) with a variety of cues (Amano & Maruyama, 2011; Kauffman *et al*., 2011; Nishijima & Maruyama, 2017). We had shown in a previous study that changes in innate behaviours using the chemoattractive odorant IAA and an aversive heat stimulus leads to loss of attraction to IAA, that lasts more than twenty hours after training, hence giving rise to LTAM formation (Dahiya *et al*, 2019). Here, we propose a novel mechanism for LTAM formation and show that the effect of training *C. elegans* with a chemoattractant and heat may lead to the release of training cue specific extracellular vesicles (EVs). Further, these EVs may then be taken up by the same or by untrained counterparts that in turn allow naïve animals to gain LTAM, resulting in impaired attraction to the chemoattractant.

*Caenorhabditis elegans* can release EVs environmentally through sensory ciliary neurons, which can act as a mode of communication. Previous studies have shown the involvement of EVs in inter-animal communication in the context of mating ((Wang *et al*, 2014) and reviewed in (Wang *et al*, 2024a)). There are multiple ciliated EV releasing neurons (EVNs) in *C. elegans.* 21 of these are male-specific neurons in the head and tail, 6 are inner labial type 2 (IL2) neurons which are present in both males and hermaphrodites (O’Hagan *et al*, 2017). It has been previously reported that CIL-7, a myristoylated protein regulates EV biogenesis and is known to be an EV cargo. The kinesin-3 protein, KLP-6, is required for the release of EVs into the environment. Both these genes, *cil-7* and *klp-6*, are expressed in the IL2 neurons of hermaphrodite *C. elegans* (Kahn-Kirby & Bargmann, 2006; Maguire *et al*, 2015; Morsci & Barr, 2011; Peden & Barr, 2005; Wang *et al*, 2021). Recent work has also shown that multiple ciliated sensory neurons pack and export ciliary membranes and excess ciliary proteins through EVs (Lobo *et al*, 2025; Razzauti & Laurent, 2021). Here we report that EVs facilitate the transfer of LTAM from trained animals to both naïve and memory-defective *C. elegans*. The cAMP response element binding protein (CREB) has been shown to be involved in memory formation across phyla, with mutants in *creb1* shown to be defective in long-term memory formation in multiple organisms ((Dash *et al*, 1990) and reviewed in (Flavell & Greenberg, 2008; Silva *et al*, 1998)). We show that *creb1/crh-1* mutant animals that show no LTAM formation in this IAA and heat-based paradigm (Dahiya *et al*., 2019), show LTAM from exposure to the extrinsic factors (likely EVs) released by trained wild-type (WT) animals.

These data incentivised us to perform LC-MS experiments to identify the chemicals that may be sent out by trained *C. elegans*. Using the LC-MS data, we go on to show that a cocktail of three chemicals sent out by *C. elegans* during the training process, likely through EVs, allows for the animals to show no attraction specifically to IAA recapitulating associative memory without a training paradigm.

This work adds to the growing body of literature describing the role of EVs in multiple processes including cell death, neurodegenerative diseases and therapeutics (Mohamed *et al*, 2025; Wang *et al*, 2025; Wiersema *et al*, 2024; Wright *et al*, 2025).

Lastly, we report that LTAM can be transferred across species of *Caenorhabditis*, where *C. elegans* and *C. briggsae* can transfer LTAM from trained animals of one species to naïve animals of the other species. Like most previous studies involving LTAM formation in *C. elegans* and across phyla, we had speculated that the mechanism of LTAM formation as a result of IAA and heat training, was intrinsic to the organism (Dahiya *et al*., 2019). Contrary to this hypothesis, our findings in this study indicate that LTAM can be transferred from one *C. elegans* to another through environmentally released extracellular vesicles (EVs), thus challenging the conventional notion of LTAM being intrinsic to an organism.

## Materials and Methods

### Strain maintenance and transgene generation

All strains were cultured on nematode growth medium (NGM) plates, inoculated with OP50 *E. coli* bacteria, and maintained under standard conditions at 22°C, following established protocols outlined in previous literature (Brenner, 1974). The N2 Bristol strain served as the standard wild-type (WT) control for comparative analyses across all experimental procedures. A synchronous population of young adult *C. elegans* was obtained using previously established protocols (Porta-de-la-Riva *et al*, 2012). The list of strains used in this study is tabulated in Supplementary Table 1 (S1) and primers used for genotyping the strains are tabulated in Supplementary Table 2 (S2). The *klp-6* rescue construct P*klp-6*::GFP::KLP-6 (gift from Maureen Barr, details in (Wang *et al*., 2014)) was microinjected in the *klp-6* mutant *C. elegans* to prepare transgenic lines as previously described (C. Mello and Fire 1995; C. C. Mello et al. 1991).

### Assay to study long-term associative memory in *C. elegans*

All assays were performed as previously described in (Dahiya *et al*., 2019) with minor modifications. Briefly, the training paradigm involves simultaneous exposure of *C. elegans* or *C. briggsae* young adults to two cues; heat and a chemoattractant (isoamyl alcohol (IAA)/ Diacetyl (depending on the experiment)), for two minutes followed by ten minutes of rest period at 22°C. This cycle is repeated five times, and the animals are then kept at 22°C for 20 hours (illustrated in Figure 1A). This leads to the formation of long-term associative memory (LTAM), seen in the form of reduced attraction towards the chemoattractant, which can last for as long as 24 hours. LTAM is quantified at 20-24 hours from the time of training, using a readout called the chemotaxis index (CI), where CI= Displacement along IAA, heptanone or diacetyl (depending on the odorant used) gradient/Total distance traversed. The concentrations of the odorants used are as previously published (Dahiya *et al*., 2019; Zhang *et al*, 2016). CI is calculated by analysing the videos of the *C. elegans* undergoing chemotaxis with respect to the chemoattractant. The videos were analysed using FIJI, and an associated plugin, Trackmate (Ershov *et al*, 2022; Schindelin *et al*, 2012). Each plot indicates a set of 3-6 replicates with all conditions/genotypes in the graph performed at the same time over multiple days. Each replicate was performed using 8-10 *C. elegans*. A detailed experimental protocol can be found in the Supplementary section.

**Figure 1:**
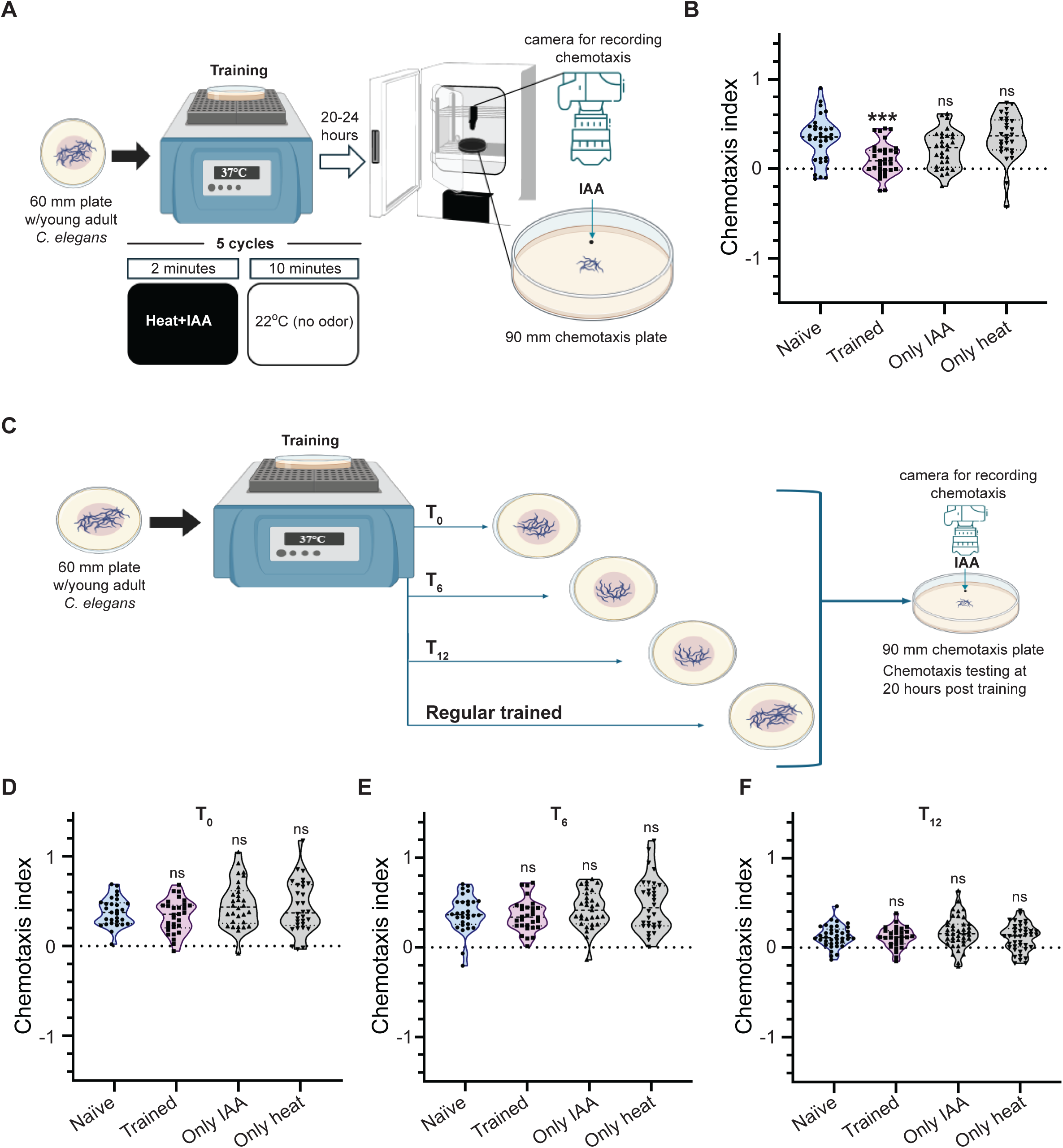
Long-term associative memory (LTAM) in *C. elegans* requires the release of extrinsic factor(s) by trained animals. (A) Illustration of the long-term associative memory (LTAM) training paradigm used in this study. (B) Violin plots showing chemotaxis index (CI) in *C. elegans*. The presence of LTAM is indicated as a reduction in the chemotaxis index (CI) of trained animals. (C) Illustration of the time-point assay to test the time of release of factor(s) required for LTAM formation. (D, E, F) Violin plots showing CI of *C. elegans* that were removed at time-points t_0_ (D), t_6_ (E) and t_12_ (F) hours respectively from plates on which the animals were trained. Multiple comparisons were performed using one-way ANOVA and p-values were adjusted using Dunnett’s correction, “***” indicates p=0.0002 and “ns” indicates not significant.

### Exchange assays

Naïve animals were transferred to an empty plate, followed by trained animals being transferred to the plate that previously had naïve *C. elegans.* The final step involved transferring the naïve animals to the plate that had had trained *C. elegans* (illustrated in Figure 2A). This exchange protocol was completed within 30 minutes from the end of training. After exchange, *C. elegans* were kept in a 22°C incubator and LTAM was tested using the chemotaxis assay 20-24 hours after training. The exchange assays were performed between *C. elegans* and *C. briggsae* in a similar manner with naïve *C. elegans* transferred to plates that had contained trained *C. briggsae* and naïve *C. briggsae* transferred to plates that had had trained *C. elegans*. All plates used for training and transfer were 60 mm in diameter.

**Figure 2:**
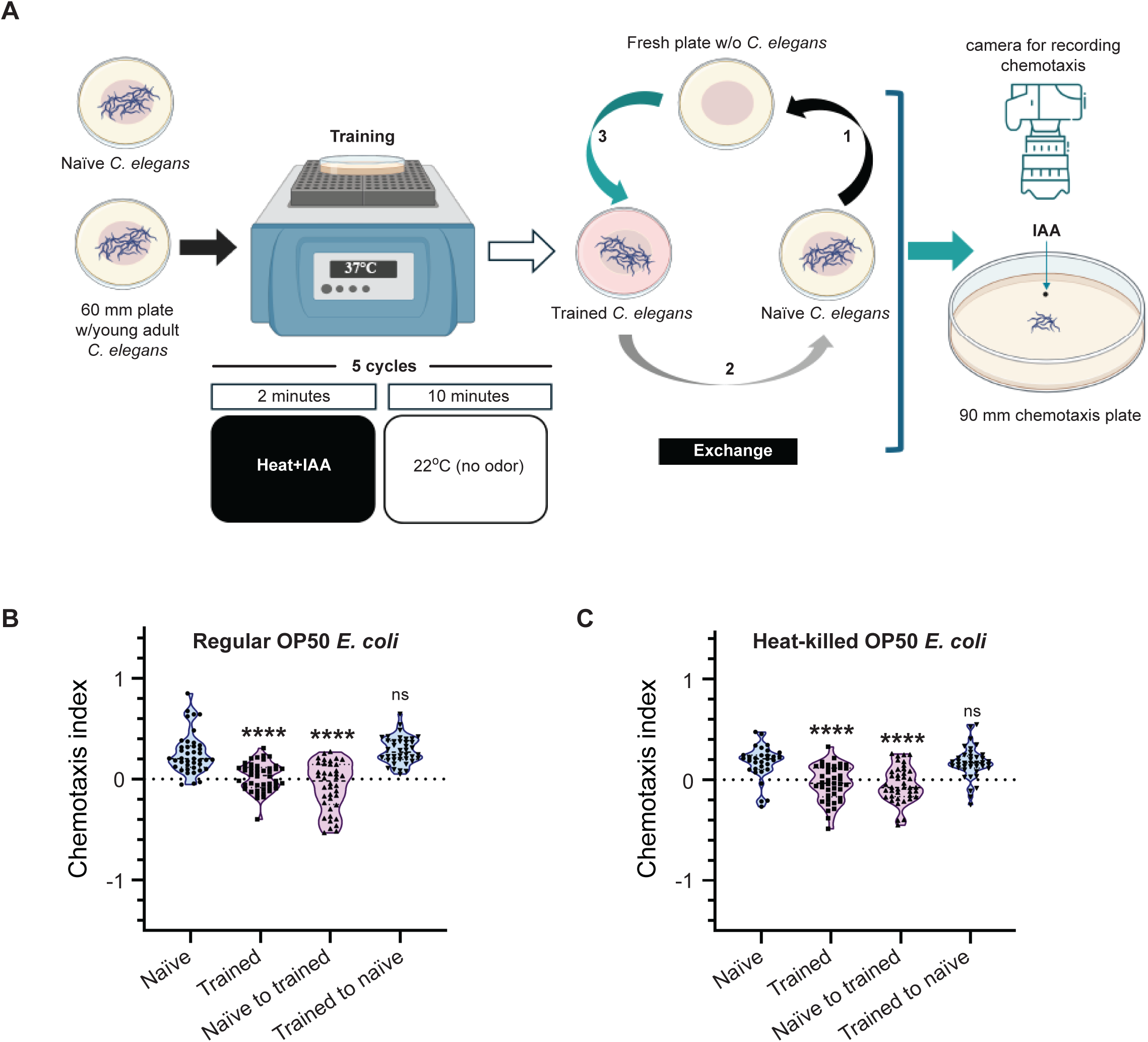
LTAM is gained by naïve *C. elegans* from trained plates. (A) Schematic of the exchange assay to expose naïve *C. elegans* to factor(s) released by freshly trained animals. The numbers along the arrows indicate the sequence of moving *C. elegans* across multiple plates. (B) Violin plots showing CI in naïve *C. elegans* when exposed to factor(s) released by trained animals. (C) Violin plots showing CI in *C. elegans* trained on plates with heat-killed bacteria. Multiple comparisons were performed using one-way ANOVA and p-values were adjusted using Dunnett’s correction, “****” indicates p< 0.0001 and “ns” indicates not significant.

### Imaging experiments

Freshly trained young adult hermaphrodite *C. elegans* were used for imaging as previously described with minor modifications (Tikiyani *et al*, 2018). Briefly, *C. elegans* were mounted on 1% agarose pads with 34 mg/ml 2,3-Butanedione Monoxime (BDM) solution in M9 medium. Images were acquired as Z-stacks on a Zeiss Apotome microscope using the 63X oil objective within four hours of the training paradigm. The order of imaging of trained and control animals was changed for each of the three replicates. For processing and analysis of images, FIJI (Schindelin *et al*., 2012) was used to quantify fluorescence intensity of cell body and number of puncta in each condition by selecting specific regions of interest. A threshold was set for analysing particles of each *C. elegans* to separately quantify cell body intensity and punctal number across the neuronal processes. The summation of mean grey values using “Analyze Particles” from each cell body per worm and the punctal number along each neuronal process was counted using “Analyze Particles”. Both cell body intensity and punctal numbers were plotted as independent graphs. Each replicate had 4-7 *C. elegans* that were imaged per condition over three replicates and each dot on the plots indicates a single animal.

### Liquid Chromatography-Mass Spectrometry (LC-MS)

The protocol used was modified from a previous study (Nikonorova *et al*, 2022). Sample collection involved four 60 mm plates with approximately 70 *C. elegans*/plate for each sample. Freshly trained WT animals were washed with M9 buffer and collected as one sample. This was repeated for every group of *C. elegans*; naïve, trained, only IAA and only heat. Next differential centrifugation was performed. Firstly, the *C. elegans* suspension was centrifuged at 3,000 g for 15 minutes at 15°C to pellet the animals and bacteria. This supernatant was transferred to fresh tubes and then centrifuged at 10,000 g for 30 minutes at 4°C. This step was repeated thrice. Lastly, the collected supernatant was sent to the IISc LC-MS facility for identification of extrinsic factor(s). Samples were similarly collected from the EV release defective mutants as naïve and trained animals as well as empty plates with and without *E. coli*. Each sample was processed in triplicate over multiple days.

### Chemical supplementation assays

The surface of unseeded NGM plates were supplemented with 300 µl of 100 mM imazapyr (Sigma Aldrich, cat. no. 37877), 2-methoxy 5-methyl aniline (2M5M, Sigma Aldrich, cat. no. 103284) or sodium salt of glycochenodeoxycholic acid (SGCDC, SRL, cat. no. 97971). For the fourth condition, all three chemicals were supplemented together (100 µl each of 100 mM of imazapyr, 2M5M and SGCDC). These plates were freshly seeded with OP50 *E. coli* bacteria after the chemical solutions had dried. Finally, a synchronized young adult population of naïve *C. elegans* was transferred (30-35 animals) to each plate. Chemotaxis to IAA and diacetyl was tested at 3.5 hours and 6 hours from the time-point of supplementation. These concentrations and the time for assays were chosen after standardization procedures involving multiple concentrations of the chemicals and multiple time-points at which the assays were performed.

### Statistical analysis

All statistical analyses were performed using GraphPad Prism 8. Outliers were identified using Grubbs method (α = 0.05) for all plots. For all plots, the mean values were compared using one-way ANOVA with Dunnett’s correction. The level of significance was set as p ≤ 0.05. The statistical analyses used for each plot is indicated in the figure legends.

## Results

### Long-term Associative memory in *C. elegans* requires extrinsic factor(s) released by the animals

We have previously published a training paradigm to study long-term associative memory (LTAM) in wild-type (WT) *C. elegans* using isoamyl alcohol (IAA) and heat ((Dahiya *et al*., 2019) and illustrated in Figure 1A). Using this paradigm, we replicated the data showing that unlike naïve animals, their trained counterparts showed a significantly lower chemotaxis index (CI) to IAA (Figure 1B, tracks indicated in Figure S1A). To investigate the underlying molecular mechanism of LTAM formation we planned to perform RNA sequencing experiments using naïve and trained WT animals.

In order to perform this experiment optimally, adult *C. elegans* had to be synchronized (without eggs and L1 animals) at 20 hours post training for RNA extraction ((Brenner, 1974) and illustrated in Figure S1B)). To remove eggs and L1s from the population, the *C. elegans* were separated from eggs twice (at 12 and 8 hours) during the twenty-hour resting stage after training and a similar protocol was followed for naïve animals (illustrated in Figure S1B). We initially gently washed the *C. elegans* and eggs with M9 to remove the *C. elegans* from the eggs on the plate at both time points. However, upon testing the animals for LTAM after the washes, we found the animals no longer showed LTAM (Figure S1C). This prompted us to pick animals at the standardized 12-hour time point during the 20-hour resting period to circumvent the effects of washing. Again, we found that the animals removed from the plates at the 12-hour time-point no longer showed LTAM (Figure 1F). This prompted us to hypothesize that trained *C. elegans* may be releasing factor(s) onto the plate that could allow for LTAM formation. In order to test the time-point at which the animals may be releasing unknown factor(s) we tested the animals by picking them from the plate within a few minutes of training (t_0_), 6 hours after training (t_6_), 12 hours after training (t_12_), and keeping some animals in the original plate (illustrated in Figure 1C). We found that only the animals on the original plate showed LTAM formation (Figure 1B), while all other animals lost LTAM associated with heat and IAA (Figures 1D-F).

Our experiments indicated that removal of *C. elegans* to new plates immediately after training led to a loss of LTAM allowing us to conclude that *C. elegans* may be releasing unknown factor(s) during training or immediately after training. We then decided to test if these factor(s) could impart LTAM to naïve animals.

### Naïve *C. elegans* on trained plates gain LTAM

To investigate if the factor(s) released by *C. elegans* during training were indeed required for LTAM formation, we decided to expose naïve *C. elegans* to the factor(s) released by the animals during training. For this, we performed an exchange assay, where naïve animals were transferred to plates that previously had trained animals, and vice versa (illustrated in Figure 2A). This exchange was performed within 30 minutes of training. Intriguingly, we observed that naïve *C. elegans* that were transferred to plates that had had trained animals showed LTAM, similar to what we observed in trained control animals (Figure 2B, tracks indicated in Figure S2A). Consistent with our previous results, trained animals that were transferred to plates that had had naïve *C. elegans*, showed a loss of LTAM (Figure 2B, tracks indicated in Figure S2A). This led us to conclude that these factor(s) released by *C. elegans* during training are sufficient to induce LTAM in naïve animals.

Next, to make sure that the factor(s) released on the plate were not produced by bacteria on the plate as the animals are well fed with OP50 *E. coli* throughout their training and resting phases, we performed the same experiments with heat-killed *E. coli.* The *E. coli* were heat-killed as previously described (Qi *et al*, 2017). We observed the same results as previously seen with live *E. coli* (Figure 2C). To confirm the associative nature of the transferred LTAM, we tested WT *C. elegans* with heat and IAA in an unpaired assay. In order to perform this experiment, we exposed naïve *C. elegans* to heat and IAA one after the other as illustrated in Figure S2B and found that in neither case could the animals form LTAM. Further, the plates that had had trained *C. elegans* did not allow for naïve *C. elegans* placed on them to gain LTAM (Figures S2C and D).

Our data so far indicates that the associative training paradigm allows *C. elegans* to release factor(s) onto its surroundings that may be taken up by naïve animals which then go on to show LTAM, hence allowing for the horizontal transfer of LTAM across animals. Next, we wanted to understand the specificity of these released factor(s).

### The training cue lends specificity to factor(s) released by trained *C. elegans*

Our results so far indicate horizontal transfer of LTAM in naïve animals when transferred to a plate that had previously contained trained animals. We wanted to understand the specificity of the released factor(s). Different odours can be sensed by largely distinct neurons in *C. elegans* (reviewed in (Chou et al, 1996)). IAA is known to be sensed by the AWC neurons, that can also sense heptanone ((Colbert & Bargmann, 1995; Zhang *et al*., 2016) and illustrated in Figure S3). In order to test the specificity of the secreted factor(s) we took animals trained using the IAA and heat paradigm, along with naïve animals transferred to plates that had contained trained *C. elegans* and tested their chemoattraction towards heptanone. We observed that these *C. elegans* showed no LTAM formation with respect to heptanone (Figure 3A). These results indicate a specificity of the released factor(s) that allow for LTAM formation only with respect to the odour used for training.

**Figure 3:**
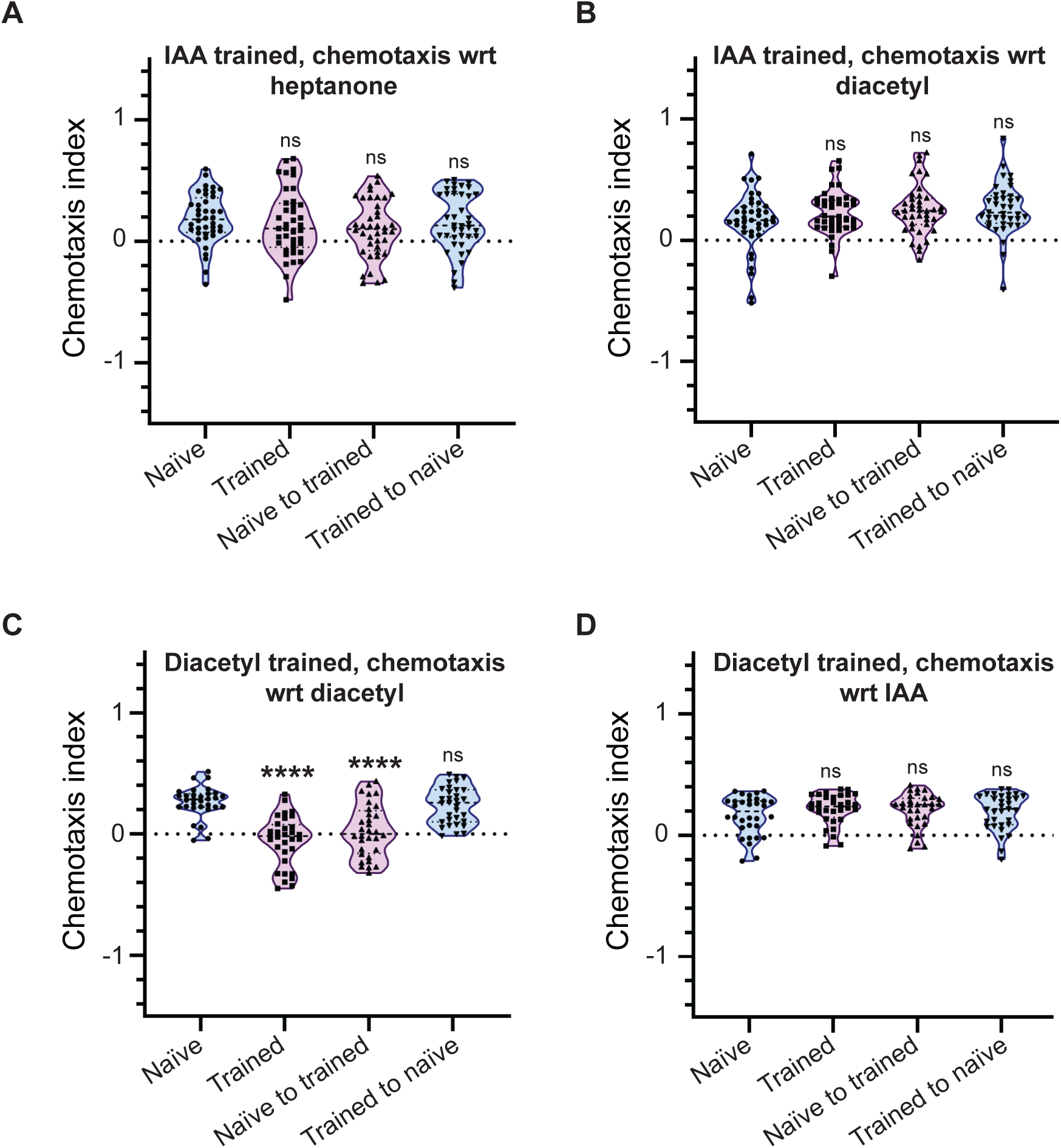
The factor(s) released during training are cue-specific. (A, B) Violin plots showing CI of animals trained using IAA and tested with heptanone (A) and diacetyl (B). Animals trained using IAA show no decrease in CI towards heptanone or diacetyl indicating no LTAM towards a chemoattractant not used during training. (C) Violin plots indicating diacetyl specific CI in diacetyl and heat trained *C. elegans* and naïve animals exposed to factor(s) released by trained animals. (D) Violin plots showing CI of animals trained with diacetyl and tested with IAA. Multiple comparisons were performed using one-way ANOVA and p-values were adjusted using Dunnett’s correction, “****” indicates p< 0.0001 and “ns” indicates not significant.

We next decided to test LTAM formation with respect to another chemoattractive odour sensed by a different neuron. Diacetyl is an odour that is sensed by the AWA neuron ((Sengupta *et al*, 1996; Sengupta *et al*, 1994) and illustrated in Figure S3). We trained animals with heat and IAA and found that trained animals and naïve animals transferred to plates that had contained trained animals did not show any defects in chemotaxis with respect to diacetyl (Figure 3B). Our previous work has shown that the heat and chemoattractant based training paradigm may be used with diacetyl and heat in a process similar to that performed with IAA and heat (Dahiya *et al*., 2019). We next trained the animals with heat and diacetyl and found that they showed LTAM towards diacetyl (Figure 3C). Further, naïve animals placed on a plate that had had trained *C. elegans* showed LTAM towards diacetyl (Figure 3C). This experiment indicates that a release of factor(s) occurs independent of whether the odour is IAA or diacetyl; the nature of these factor(s), however, appears to be extremely specific to the odorant used during training. Finally, we trained animals with diacetyl and tested them with IAA and found no LTAM formation towards IAA (Figures 3D). These data together indicate a high degree of specificity of the factor(s) released for allowing LTAM formation only with respect to the cue present in the training paradigm.

We next wanted to study the role of these factor(s) in animals that are known to be defective in memory formation.

### LTAM defective *C. elegans* show LTAM from exposure to plates that had contained trained WT animals

A plethora of literature has implicated CREB1 (cAMP Response Element Binding protein 1) in long-term memory formation across phyla ((Bartsch *et al*, 1998; Timbers & Rankin, 2011; Yin *et al*, 1995; Yin *et al*, 1994) and reviewed in (Flavell & Greenberg, 2008; Johannessen *et al*, 2004; Silva *et al*., 1998; Yin & Tully, 1996)). We have shown that the mutants of the ortholog of mammalian *creb1*, *crh-1* in *C. elegans,* show LTAM defects in the heat and IAA paired assay (Dahiya *et al*., 2019). We decided to test if *crh-1* mutants allowed for LTAM formation when transferred to a plate that had contained trained WT animals (illustrated in Figure 4A). Our data indicates that *crh-1* mutants that basally show no LTAM, show LTAM when transferred to a plate which had had WT trained animals. We also found that WT trained animals transferred to a plate that had contained trained *crh-1* mutant *C. elegans* do not show LTAM (Figure 4B and tracks indicated in Figure S4). These data allowed us to conclude that *crh-1* mutants may have defects in the synthesis or release of the factor(s) that allow for LTAM formation but appear to be able to uptake these factor(s) to show LTAM.

**Figure 4:**
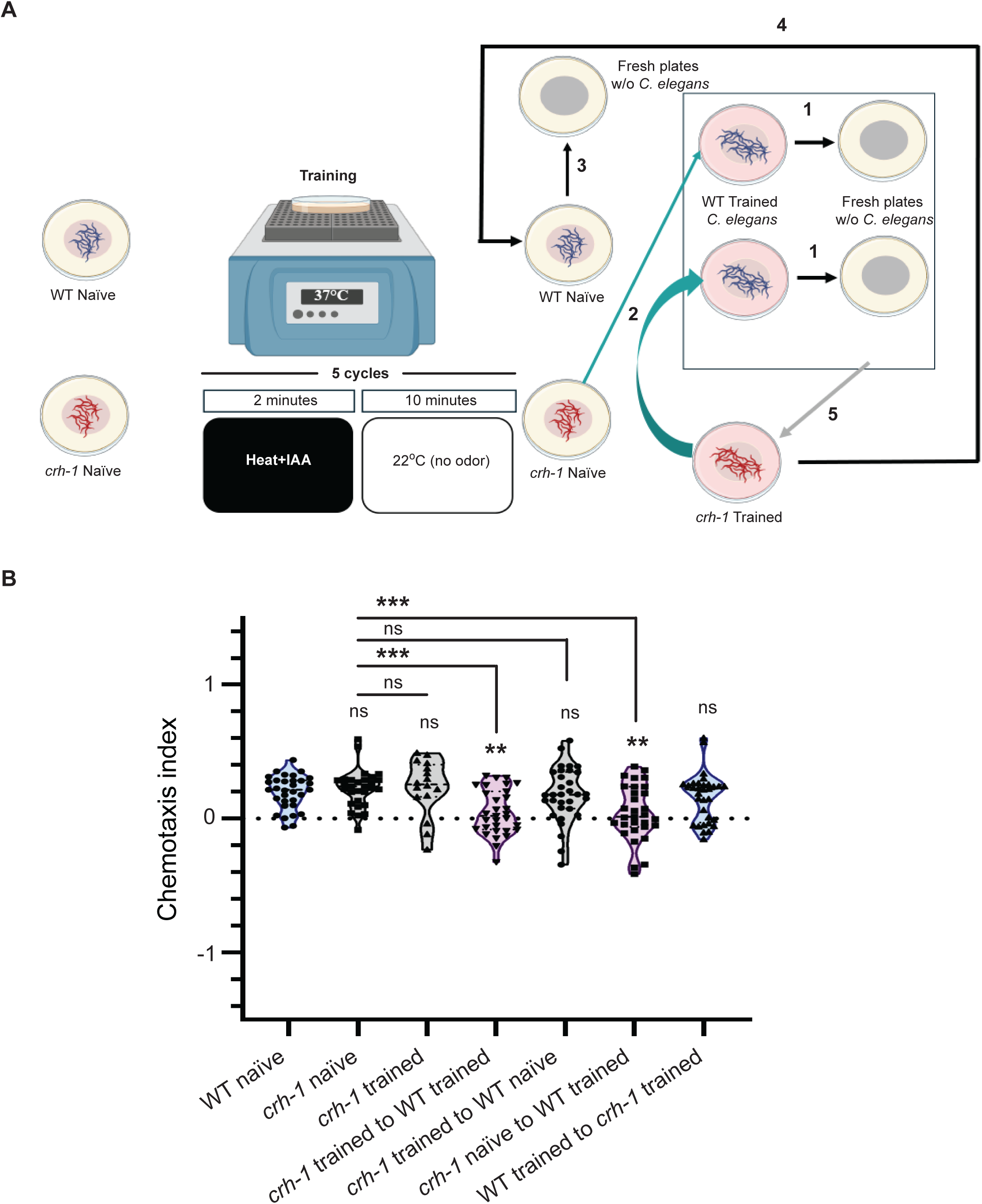
LTAM can be gained by memory defective *crh-1* mutant *C. elegans* from trained plates that contained wild-type animals. (A) Schematic representation of exchange assay between wild-type (WT) and *crh-1* mutant *C. elegans.* The numbers along the arrows indicate the sequence of moving *C. elegans* across multiple plates. (B) Violin plots indicating CI in *crh-1* mutant animals (naïve and trained) when exposed to factor(s) released by trained WT animals. Multiple comparisons were performed using one-way ANOVA and p-values were adjusted using Dunnett’s correction and Sidak’s correction (for within group comparisons), “**” indicates p< 0.01, “***” indicates p< 0.001 and “ns” indicates not significant. Note that in the *crh-1* trained violin plot, only 16 animals are plotted across 3 replicates, although 8 *C. elegans* were taken per replicate. This is because of multiple tracks that overlapped during analyses and could not be separated out.

We next wanted to gain insight into the mechanism of secretion of the factor(s) during training of *C. elegans*.

### Extracellular vesicles are required for LTAM formation

A well-studied mechanism of social behaviour in *C. elegans* is through secreted small molecule signalling by ascarosides (reviewed in (Yang *et al*, 2023)). Pre-exposure to different ascarosides can change the chemotaxis of *C. elegans* to different odours (Wu *et al*, 2023). We decided to test for LTAM formation in the *daf-22* mutant *C. elegans* that are defective in ascaroside biosynthesis (Golden & Riddle, 1985). We found that *daf-22* mutants showed no defects in LTAM formation (Figure 5A). These data indicate that the secreted factor(s) may not be *daf-22* dependent. Apart from ascarosides, sensory signalling allowing for behaviours like mate sensing through chemoattraction requires extracellular vesicle (EV) release from *C. elegans* (reviewed in (Wang *et al*., 2024a)). We next tested mutants in EV biogenesis (*cil-7*) and release (*klp-6*) for LTAM formation (Wang *et al*., 2021). We found that both mutants showed no LTAM formation (Figure 5B and C). In order to further confirm the role of EVs in the process of LTAM, we attempted to rescue the loss of LTAM phenotype in *klp-6* mutants by expressing KLP-6 specifically in the IL2 neurons where KLP-6 has been shown to be expressed in hermaphrodites (Peden & Barr, 2005; Wang *et al*, 2024b). We observed that expressing KLP-6 in the mutant *klp-6* animals rescued the loss of LTAM phenotype where both trained *C. elegans* as well as naïve animals transferred to plates that had contained trained animals now showed LTAM formation (Figure 5D). We next decided to test if *klp-6* mutants allowed for LTAM formation when transferred to a plate that had trained WT animals. We found that trained *klp-6* mutants transferred to a plates which had had trained WT animals showed LTAM, while trained WT animals transferred to plates that had contained trained *klp-6* mutant *C. elegans* did not show LTAM (Figure 5E and tracks indicated in Figure S5). These data allow us to conclude that EV release may be the basis for LTAM formation in a chemoattractant and heat-based associative training paradigm.

**Figure 5:**
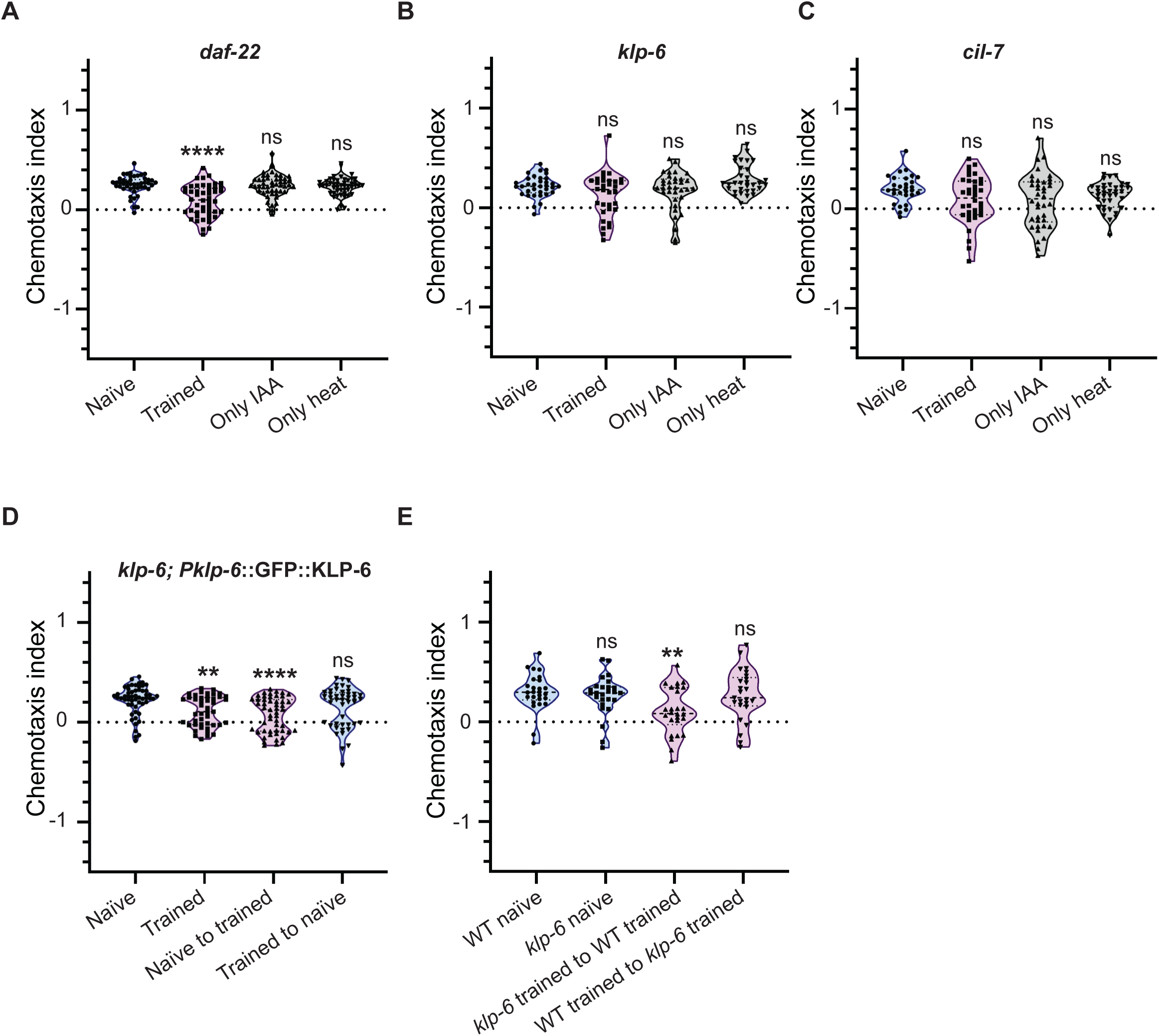
Role of extracellular vesicles released from IL2 neurons in LTAM formation and transfer. (A) Violin plots indicating CI in *daf-22* mutant animals defective in ascaroside biosynthesis. (B) Violin plots indicating CI in *klp-6* mutant animals defective in extracellular vesicle release. These animals show no LTAM formation. (C) Violin plots indicating CI in *cil-7* mutant animals defective in extracellular vesicle biosynthesis. These animals also show no LTAM formation. (D) Violin plots indicating CI in the *klp-6* rescuing strain. (E) Violin plots indicating CI in *klp-6* mutant animals when exposed to factor(s) released by WT animals, no LTAM was observed in WT animals exposed to plates that had had trained *klp-6* mutants. Multiple comparisons for all plots were performed using one-way ANOVA and p-values were adjusted using Dunnett’s correction, “**” indicates p< 0.01, “****” indicates p< 0.0001 and “ns” indicates not significant.

We next investigated changes in EV expression in the nervous system upon training *C. elegans* with heat and IAA.

### Changes in EV biogenesis marker expression upon heat and IAA training

Previous work has elegantly shown that an EV cargo, CIL-7, is released from the IL2 neurons into the external region around *C. elegans* (Wang *et al*., 2024b). We tested the same line for changes in fluorescence expression in the IL2 neurons and found that the *cil-7 (my61[cil-7::mNG])I; him-5(e1490)* line showed increased fluorescence in the IL2 cell bodies and an increase in fluorescent puncta along the IL2 neuronal processes in trained WT animals when compared to the control *C. elegans* that were either naïve or trained with only IAA or only heat (Figures 6A and B). With these data indicating the involvement of EVs in LTAM transfer. We next decided to perform a Liquid Chromatography-Mass Spectrometry (LC-MS) to study the differences in the released products upon heat and IAA training when compared to control animals.

**Figure 6:**
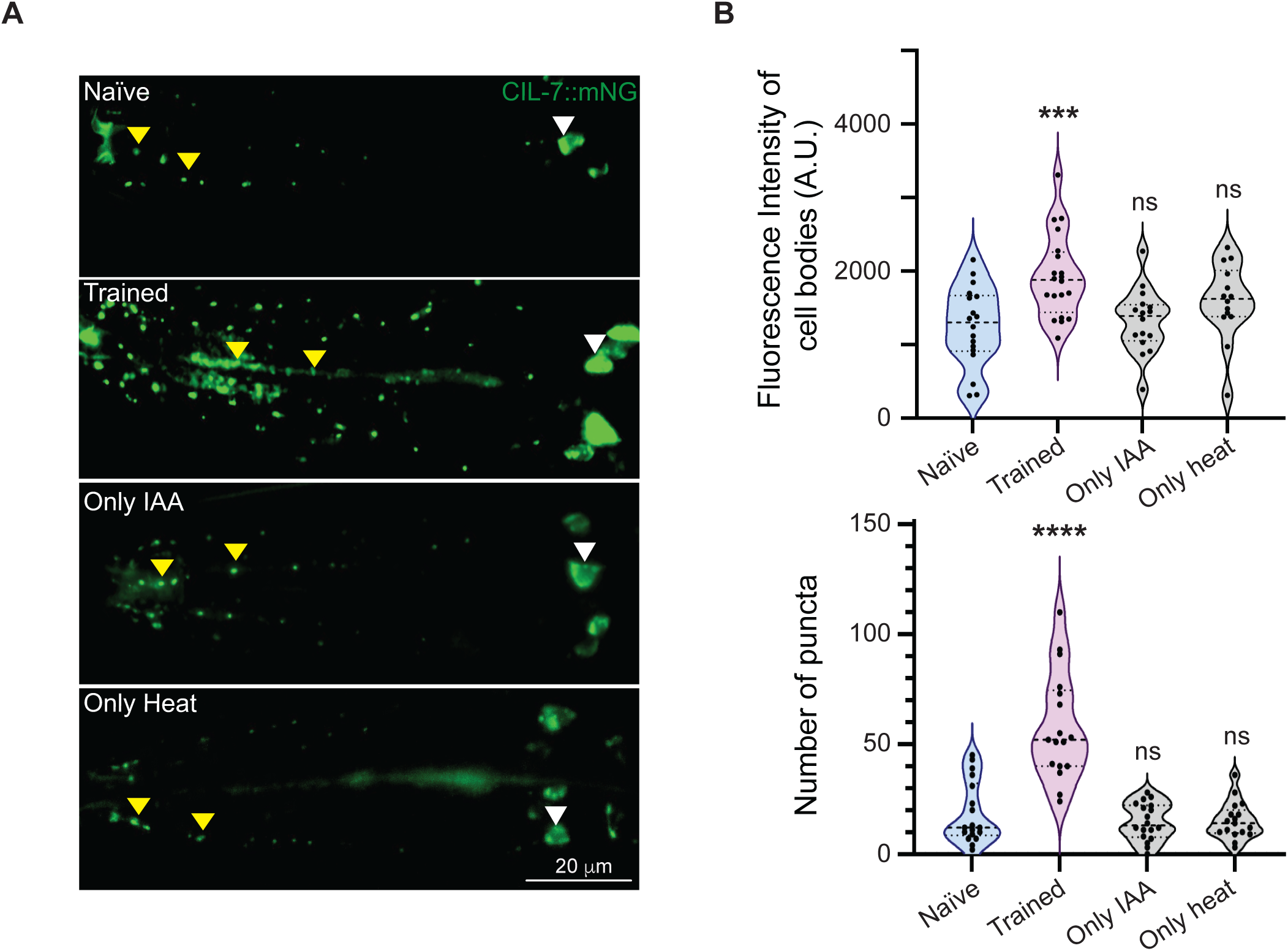
Increase in EVs in trained *C. elegans*. (A) Representative fluorescence imaging of the IL2 neurons using the *cil-7*::mNG marker line in naïve, trained, IAA only and heat only conditions. (B) Violin plots showing total fluorescence intensity of cell bodies of IL2 neurons (above) and the number of puncta across the IL2 processes (below). The trained animals show increased cell body fluorescence and increased punctal number along the IL2 processes when compared to the control *C. elegans*. White arrowheads indicate cell bodies and yellow arrowheads indicate puncta in all images. Multiple comparisons were performed using one-way ANOVA and p-values were adjusted using Dunnett’s correction, “***” indicates p< 0.001, “****” indicates p< 0.0001 and “ns” indicates not significant.

### A mixture of three chemicals mimics memory formation in *C. elegans*

LC-MS was performed using WT *C. elegans* as well as *klp-6* mutant animals. Multiple LC-MS experiments were performed, each in triplicate as indicated in Figure 7A. This experiment allowed us to find 9 consistent hits that were only present in the heat and IAA trained *C. elegans* plates (Figure 7A and S6). These data suggest that multiple ions are differentially released onto the plates upon heat and IAA training. These ions may allow for LTAM formation in *C. elegans*. Based on the availability of chemicals, we supplemented the plates of naïve animals with three chemicals from our candidate list – imazapyr, 2M5M and SGCDC. Each molecule was supplemented on NGM plates onto which naïve animals were placed. To understand their cumulative role, we also supplemented all three chemicals together.

**Figure 7:**
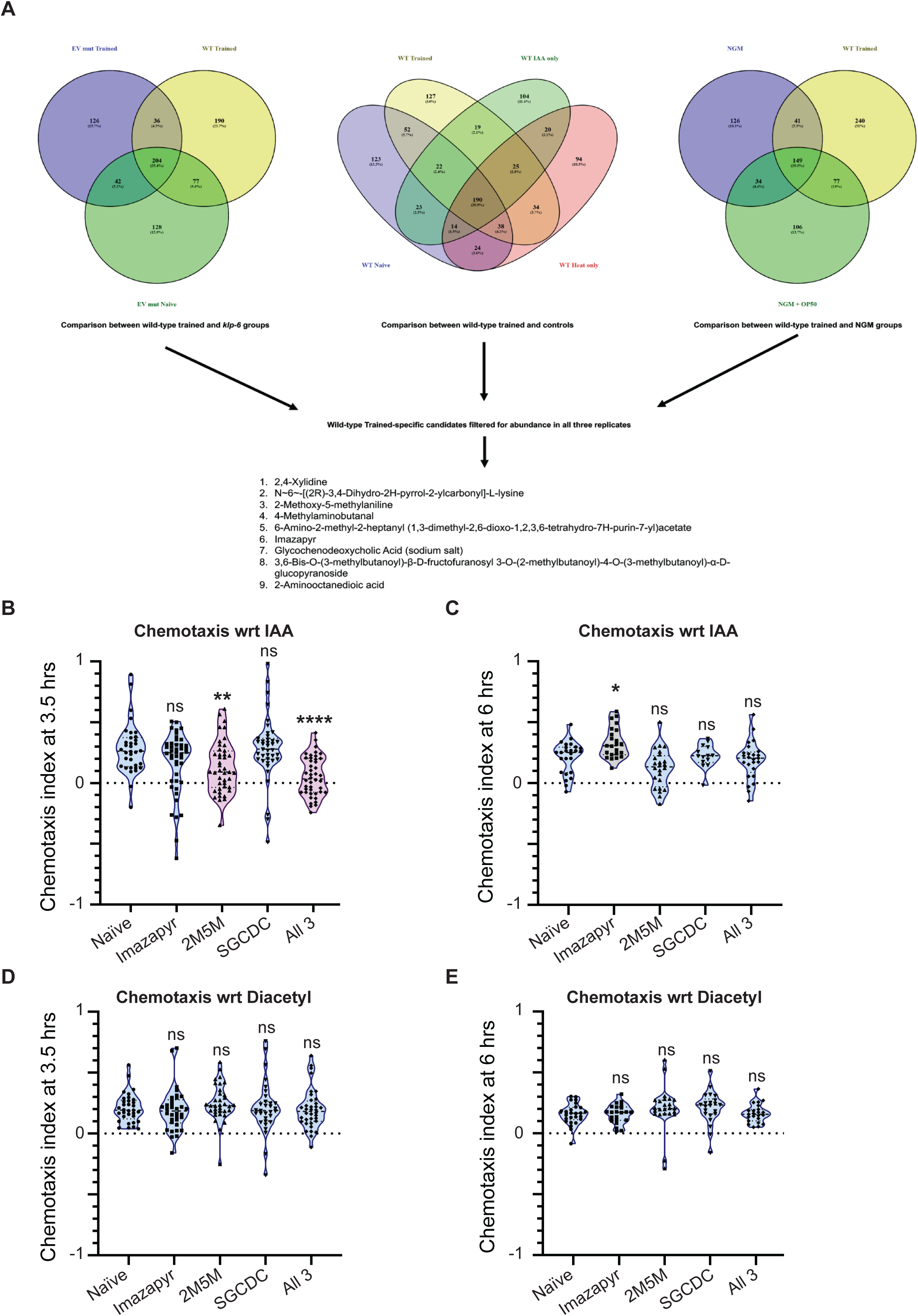
A cocktail of molecules released during training from C. elegans is sufficient to induce memory formation in *C. elegans*. (A) LC-MS data analysed from eight animal groups processed as three replicates per group; WT trained and controls (4 groups), *klp-6* naïve and trained *C. elegans* (2 groups) and empty plates containing NGM alone or NGM+OP50 *E. coli* (2 groups). The venn diagrams were created using the Venny 2.1 software (https://bioinfogp.cnb.csic.es/tools/venny/). The list of molecules present exclusively on WT trained plates in every replicate is indicated at in the bottom panel of 7A. (B) Violin plots indicating CI towards IAA in naïve *C. elegans* placed on NGM plates with OP50 containing nothing (naïve), Imazapyr, 2M5M, SGCDC or all 3 chemicals. This experiment was performed at 3.5 hours after placing young adult animals on the plate. (C) Violin plots indicating CI towards IAA in naïve *C. elegans* placed on NGM plates with OP50 containing nothing (naïve), Imazapyr, 2M5M, SGCDC or all 3 chemicals. This experiment was performed at 6 hours after placing young adult animals on the plate. (D) Violin plots indicating CI towards diacetyl in naïve *C. elegans* placed on NGM plates with OP50 containing nothing (naïve), Imazapyr, 2M5M, SGCDC or all 3 chemicals. This experiment was performed at 3.5 hours after placing young adult animals on the plate. (E) Violin plots indicating CI towards diacetyl in naïve *C. elegans* placed on NGM plates with OP50 containing nothing (naïve), Imazapyr, 2M5M, SGCDC or all 3 chemicals. This experiment was performed at 6 hours after placing young adult animals on the plate. Multiple comparisons were performed using one-way ANOVA and p-values were adjusted using Dunnett’s correction and Sidak’s correction (for within group comparisons), “*” indicates p< 0.05, “**” indicates p< 0.01, “****” indicates p< 0.0001 and “ns” indicates not significant.

We observed that imazapyr or SGCDC alone did not alter the chemotaxis towards IAA in treated *C. elegans*, whereas 2M5M supplementation led to the loss of attraction to IAA, seen as a lower chemotaxis index when compared to naïve animals This loss of attraction to IAA phenotype was more pronounced in *C. elegans* treated with all three molecules (Figure 7B, and track images in S7A). This effect was seen at the 3.5 hours’ time-point from supplementation and is lost by 6 hours after supplementation (Figure 7C), indicating that these molecules allow for memory formation, although not LTAM formation. This could be because of the absence of the other six molecules from our candidate list, along with other cargo including nucleic acids that are usually known to be carried by EVs released into the environment (Nikonorova *et al*., 2022). We also checked the chemotaxis of naïve and animals trained with IAA and heat towards Diacetyl at the 3.5- and 6-hour time-points.

At both time points we observed no loss of attraction towards Diacetyl (Figures 7D and E). As controls we performed the training and testing protocols with WT *C. elegans* where testing was performed at 3.5 and 6 hours after training. The aversion to IAA was consistently seen at both time points (Figures S7B and C). These results establish our findings that specific factors released in response to training, could lead to specific loss of attraction towards the odorant used in the training paradigm.

Finally, we wanted to test if LTAM formation is conserved across *Caenorhabditis* species. In order to perform this experiment, we investigated LTAM formation in *Caenorhabditis briggsae*.

### *Caenorhabditis* show interspecies transfer of LTAM

We first tested the *C. briggsae* strain for LTAM formation to heat and IAA and found that *C. briggsae* were surprisingly not attracted to IAA even in naïve conditions (Figure S8A and B). This precluded us from studying LTAM formation in *C. briggsae* using the IAA and heat based associative paradigm. However, since *C. briggsae* naïve animals behaved like *C. elegans* trained animals (Figure S8B), we went on to transfer *C. briggsae* naïve animals to a plate that had contained *C. elegans* trained animals and found that there was no change in *C. briggsae* chemotaxis index. We also transferred trained *C. elegans* to a plate that had contained naïve *C. briggsae* (exchange assay) and found that these animals behaved like naïve *C. elegans* (Figure S8B). These data could not allow us to make any conclusions other than the fact that *C. briggsae* was not attracted to IAA in the naïve condition unlike *C. elegans*.

We next tested if *C. briggsae* was attracted to diacetyl. We found that *C. briggsae* was both attracted to and formed LTAM with heat and diacetyl (Figure 8A). We went on to transfer *C. briggsae* naïve animals to a plate that had contained *C. briggsae* trained animals and vice versa. We found that *C. briggsae* naïve animals acquired LTAM from the plate that had contained trained *C. briggsae*, and surprisingly the trained *C. briggsae* did not lose LTAM from being on a plate that had had naïve *C. briggsae* (Figure 8B). These data indicate that a similar mechanism may allow for LTAM transfer in *C. briggsae* as is seen in *C. elegans*, however the LTAM appears to be more stable in *C. briggsae* when compared to *C. elegans*. Finally, we tested if interspecies transfer of LTAM could occur. For this experiment we transferred naïve *C. briggsae* to a plate that had contained trained *C. elegans* and transferred naïve *C. elegans* to a plate that had contained trained *C. briggsae* and found that both sets of animals showed LTAM (Figure 8C). We also transferred trained *C. elegans* to a plate that had contained naïve *C. briggsae* and unsurprisingly found no LTAM formation (Figure 8C).

**Figure 8:**
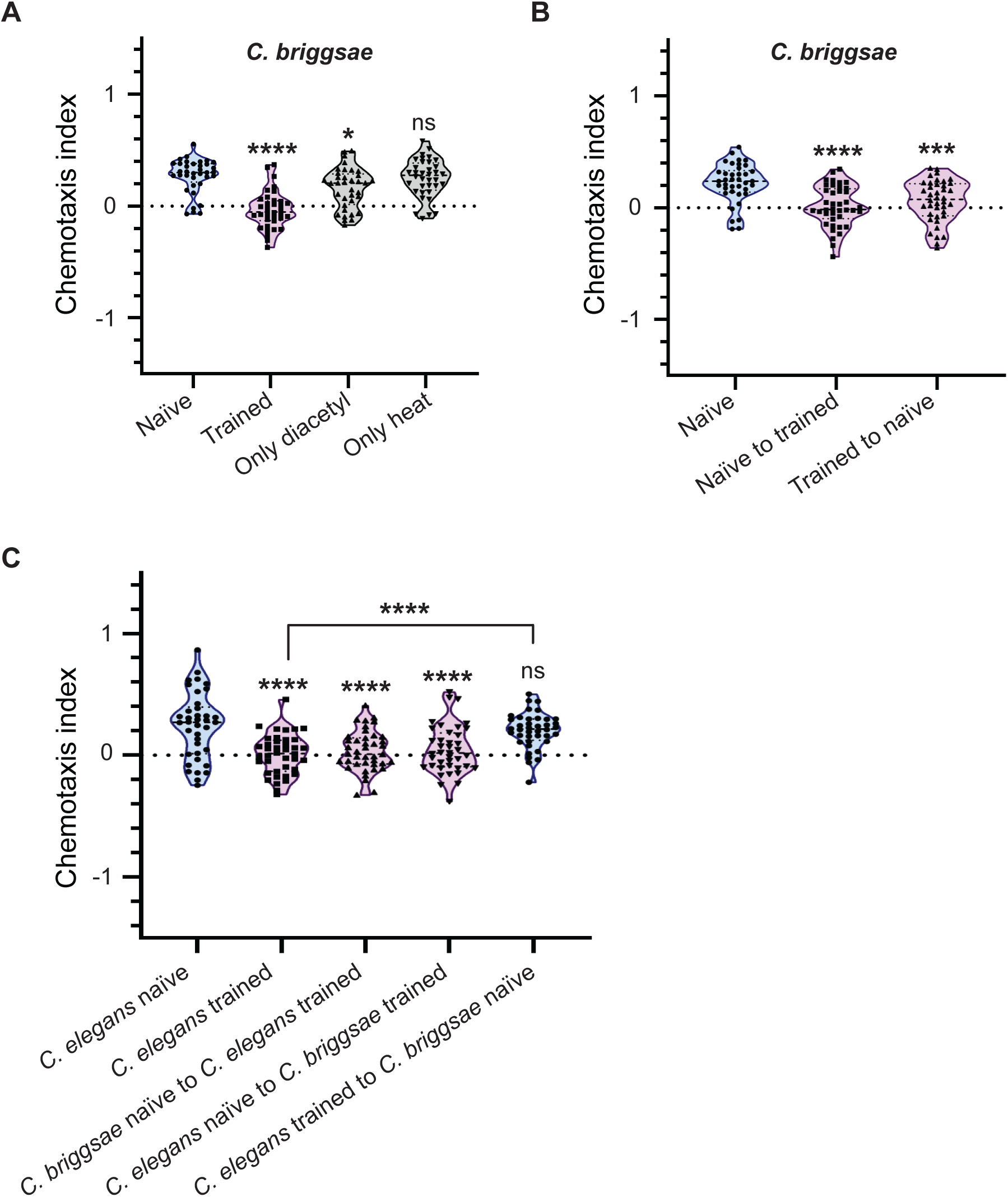
LTAM formation and transfer is conserved across *Caenorhabditis* species. (A) Violin plots indicating CI in *C. briggsae* trained with the Diacetyl and heat associative learning paradigm. (B) Violin plots indicating CI in *C. briggsae* that underwent exchange from trained plates. This plot indicates conservation of the LTAM transfer phenomenon from naïve to trained animals in *C. briggsae*. Further, no LTAM loss is seen in trained *C. briggsae* when removed from trained plates, unlike in *C. elegans*. (C) Violin plots indicating CI after transferring *C. briggsae* to plates that had contained trained *C. elegans* and transferring *C. elegans* to plates that had contained trained *C. briggsae*. This plot indicates interspecies transfer of LTAM from *C. elegans* to *C. briggsae* and vice versa. Multiple comparisons were performed using one-way ANOVA and p-values were adjusted using Dunnett’s correction and Sidak’s correction (for within group comparisons), “*” indicates p< 0.05, “***” indicates p< 0.001, “****” indicates p< 0.0001 and “ns” indicates not significant.

Together, our data indicates the role of extracellular vesicles in LTAM formation and its horizontal transfer in *C. elegans* along with its possible role in this phenomenon across species.

## Discussion

In this study we report three novel findings that have broad implications. First, *C. elegans* form long-term associative memory (LTAM) through environmentally released extracellular vesicles (EVs) from the IL2 neurons when the animals undergo training with heat and a chemoattractant. Further, these EVs appear to be cue specific and allow for LTAM formation only with respect to the cue used in the initial training paradigm. Second, LTAM can be transferred to naïve as well as memory-defective mutant animals possibly through EVs. Third, there appears to be conservation of this behaviour across *Caenorhabditis* species with EVs from *C. elegans* being able to elicit LTAM formation in *C. briggsae* and vice versa. These findings are illustrated in Figure 9. Below, we discuss the significance of these findings.

**Figure 9:**
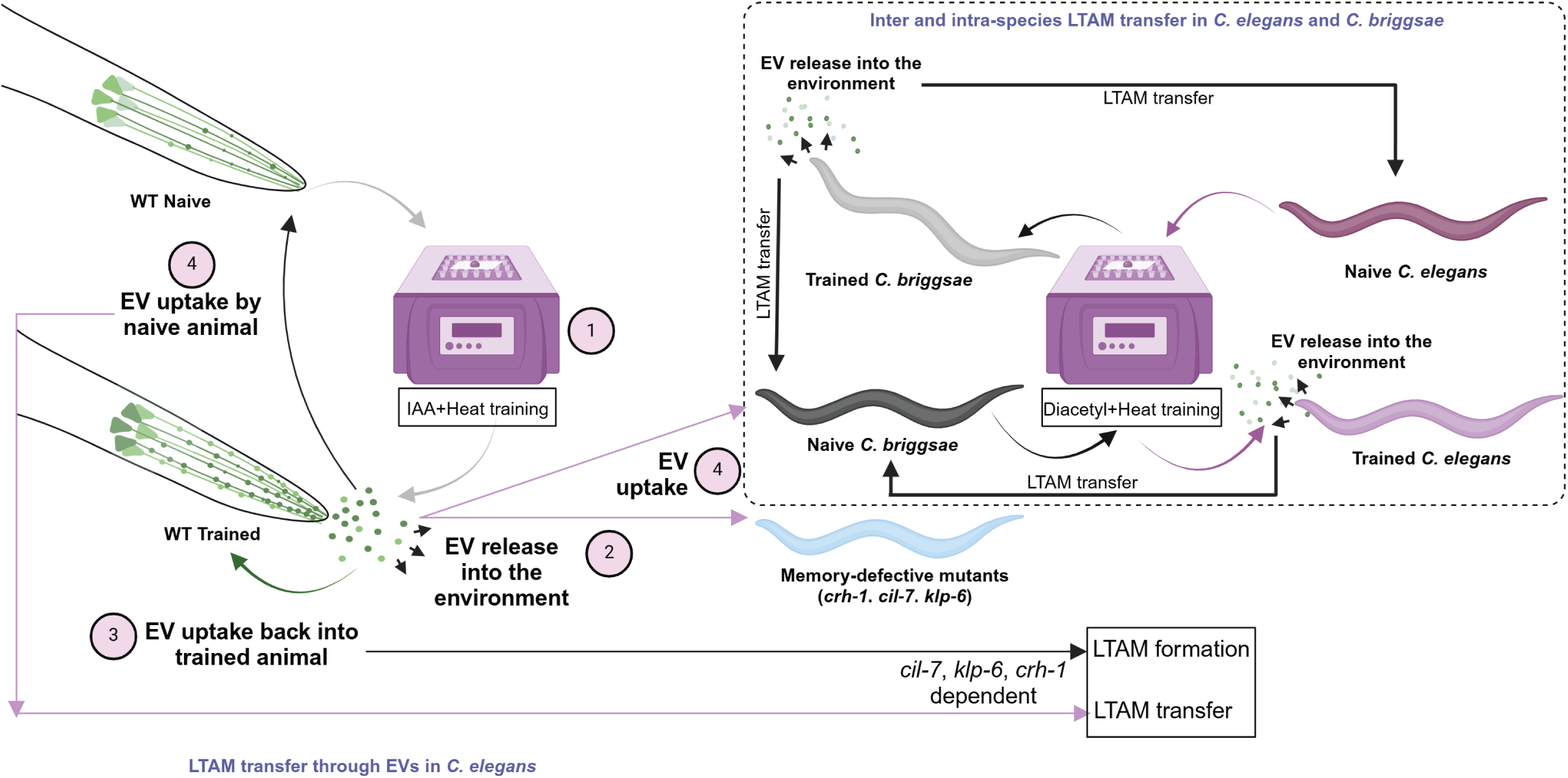
Model. Illustration of the proposed mechanistic model showing that the release of EVs from IL2 in trained *C. elegans* allows for LTAM formation in naïve animals, memory defective *crh-1* mutant *C. elegans* and EV biogenesis defective *klp-6* mutant *C. elegans*. Further, this transfer of LTAM is conserved across *Caenorhabditis*, with LTAM formation occurring in naïve *C. briggsae* from EVs of trained *C. elegans* and vice versa. The numbers indicate the order of processes including EV release followed by uptake.

### Novel mechanism for horizontal transfer of LTAM

Previous studies have shown that a form of non-associative memory can be transferred from one *Aplysia* to another by injecting RNA extracted from the trained animal into the naïve counterpart (Bedecarrats *et al*, 2018). More recent studies have shown that learned avoidance to pathogenic bacteria can be transferred from trained to naïve *C. elegans* through the *Cer1* retrotransposon (Moore *et al*, 2021). However, there appear to be no studies showing the organic transfer of LTAM from one organism to another without active communication between the animals. In this study, we show that *C. elegans* when trained using an IAA and heat-based associative learning paradigm may release extrinsic factor(s). Removal of trained animals from these plates results in loss of LTAM, indicating that these factor(s) are responsible for the process of LTAM formation. Our results also show that these factor(s) are likely released during training. Naïve animals exposed to the factor(s) released by trained animals also show LTAM formation, similar to that seen in the original trained animals. This work uncovers a novel mechanism for the horizontal transfer of LTAM through secreted EVs from one *Caenorhabditis elegans* to another, irrespective of whether the animals have been trained with the associative learning paradigm. Further, our work suggests that EVs released due to training appear to carry extremely specific cargo with respect to the odorant cue used during training. The cargo may dynamically change with changing training cues allowing for cue specific LTAM formation.

### LTAM formation in memory defective animals

Apart from allowing for LTAM formation in naïve animals, we show that the factor(s) (EVs) secreted by wild-type (WT) *C. elegans* also show LTAM in memory defective *creb1/crh-1* and EV biogenesis/cargo defective animals using multiple previously described mutants (Dahiya *et al*., 2019; Maguire *et al*., 2015; Wang *et al*., 2021; Wang *et al*., 2024b). These animals gain LTAM to the specific isoamyl alcohol (IAA) cue by being placed on a plate that had had WT animals trained with heat and IAA. This intriguing finding may allow for a better understanding of the factor(s) released and the mechanism of EV uptake/action that appear to be intact in these mutants that show defects in LTAM formation. Future work may allow for a better understanding of the involvement of CREB/CRH-1 in the observed LTAM formation and transfer phenomenon.

### A chemical cocktail that allows *C. elegans* to behave like it has been trained with heat and IAA

We found that a cocktail of Imazapyr, 2-methoxy 5-methylaniline (2M5M) and Glycochenodeoxycholic acid (SGCDC) elicits a loss of attraction specifically to IAA, albeit for a short time-span of upto 3.5 hours. How these chemicals may act throws open interesting future experiments to understand memory and chemosensation in *Caenorhabditis*. Imazapyr is known to be a non-selective herbicide which controls vegetation, especially hard-to-control perennial grass by inhibiting the acetolactate synthase (ALS) enzyme. It can disrupt protein synthesis and interfere with the synthesis of DNA and cell growth (Dersch *et al*, 2016). ALS is the first enzyme in the pathway for the synthesis of branched-chain amino acids (Dezfulian *et al*, 2017). 2M5M is an organic compound that is largely used in the production of dyes and pigments of inks and paints. It is also used as a catalyst in reactions like nitration of aromatic compounds. (https://chem-iso.com/18630). Finally, SGCDC, usually found as a sodium salt is a bile salt found in the liver, generated from chenodeoxycholic acid and glycine. It has been shown to inhibit autophagosome formation lysosomal function leading to apoptosis of human hepatocyte cells (Lan *et al*, 2020). However, to our knowledge, none of these compounds have been studied in the context of *C. elegans*. It would be interesting to test more of the compounds found in our LC-MS experiments to try and get a better understanding of what *C. elegans* release during heat and IAA training.

### LTAM transfer between Caenorhabditis briggsae and Caenorhabditis elegans

We show that *C. briggsae*, a relative of *C. elegans*, with a divergence of tens of millions of years (Butler *et al*, 1981) also show LTAM formation through released factor(s) that are likely to be EV dependent. We found that *C. briggsae* do not appear to be attracted to IAA in a manner similar to *C. elegans*, likely due to the high divergence in the olfactory pathways of these animals (Jovelin *et al*, 2009). However, both animals showed attraction towards diacetyl. We show that LTAM can transfer from trained *C. briggsae* to naïve *C. elegans* and trained *C. elegans* to naïve *C. briggsae*, indicating a conserved mechanism of LTAM transfer across *Caenorhabditis,* likely through EVs.

The one interesting difference we found was that unlike *C. elegans*, *C. briggsae* does not seem to lose LTAM when removed from the plates that they were trained on. This could be because of two possibilities; *C. briggsae* release the extrinsic factor(s) continuously for a period of time, long enough for LTAM consolidation, or these factor(s) (EVs) are stable and functional for longer periods of time in *C. briggsae.* Further studies may allow for parsing out these differences between the two species.

Our work leads to multiple outstanding questions that may allow for a better understanding of the process of LTAM transfer across organisms. We briefly discuss some of these unanswered questions below.

### Neuronal circuitry allowing for EV release by IL2 neurons for LTAM formation

Studies have shown that CIL-7, a myristoylated protein regulates EV biogenesis along with being an EV cargo and the kinesin-3 protein KLP-6 is required for the release of EVs. Both these genes, *cil-7* and *klp-6,* are expressed in the IL2 neurons in *C. elegans* hermaphrodites (Kahn-Kirby & Bargmann, 2006; Maguire *et al*., 2015; Morsci & Barr, 2011; Peden & Barr, 2005; Wang *et al*., 2021). Here we report that *cil-7* and *klp-6* mutant animals are LTAM defective, uncovering a previously unreported role of the IL2 neurons in LTAM formation.

This raises important questions in understanding the neuronal circuitry involved. Different neurons are responsible for sensing different odorants in *C. elegans*. Our experiments indicate that animals trained with IAA (sensed by the AWC neurons) and heat versus diacetyl (sensed by the AWA neurons) and heat release EVs with different cargo causing cue-specific LTAM formation. Understanding the neuronal circuitry between chemosensory neurons and IL2 neurons may allow for a better understanding of the specificity of LTAM formation.

### Dynamic EV cargo for cue-specific LTAM

*C. elegans* show LTAM formation and transfer, specific to the odorant cue used during training. For examining the extrinsic factor(s) released by trained animals, we performed LC-MS to identify candidate molecules present exclusively on the plates of trained animals (Figure 6B). One of these molecules 4-methylaminobutanal, might be of potential interest. IAA is chemically 3-methylbutanol, and a biological pathway for the conversion of 3-methylbutanol to 4-methylaminobutanal may exist. The steps would include oxidation of 3-methylbutanol to 3-methylbutanal through an alcohol dehydrogenase (Illikoud *et al*, 2018), followed by a transamination step producing 3-methylbutylamine (Encyclopedia of Sensors and Biosensors, 2023). This intermediate could potentially result in 4-methylaminobutanal through additional reactions. We have shown that a cocktail of three of these LC-MS hits allows for C. elegans to lose attraction to IAA, how these and other molecules function would allow for more insight into what EV cargoes may consist of as well allow for a better understanding of memory towards IAA. Since we see specificity of LTAM formation to the odorant used in the training paradigm, it is plausible that different odorants may allow for release of cue-specific cargo. This in turn may give rise to the specificity of the observed LTAM phenomenon.

### Uptake of EV cargo for LTAM transfer and a role for RNAs in this process

Transfer of LTAM from one *C. elegans* to another entails factor(s) (EVs) released by trained animals, and its uptake by naïve animals. This uptake could occur through a variety of mechanisms. Is the cargo released through EVs retained within the EV structure after release into the environment? Are the individual components of vesicles dispersed onto the plates? Further, does the uptake mechanism involve only neurons/glia or could other tissues like cuticles/hypodermis be involved? Understanding these aspects of the uptake mechanism may shed more light on the process of LTAM transfer across *Caenorhabditis* species.

RNA has been implicated in memory in multiple model organisms. The role of different types of non-coding RNAs like micro-RNAs (miRNAs) and Piwi-interacting RNAs (piRNAs) have been explored and studied in the context of long-term non-associative memory in *Aplysia* (Fiumara *et al*, 2015; Rajasethupathy *et al*, 2012; Rajasethupathy *et al*, 2009). Further, it has been shown in *Aplysia* that memory formed from long-term sensitization (non-associative) can be transferred from trained to naïve animals (Bedecarrats *et al*., 2018). More recently, multiple studies in have defined the role of non-coding RNAs in transgenerational memory as well (reviewed in (Miska & Rechavi, 2021)). Studies have also shown that small non-coding RNAs expressed in *Pseudomonas* strains allow for the transgenerational inheritance of avoidance memory in *C. elegans* (Kaletsky *et al*, 2020; Legue *et al*, 2022; Sengupta *et al*, 2024; Seto *et al*, 2024). Additionally, Moore et al, 2021 has shown that this avoidance memory can also be horizontally transferred from one animal to another (trained to naïve *C. elegans*) through virus-like particles encoded by the *Cer1* retrotransposon (Moore *et al*., 2021). Multiple studies have shown that extracellular vesicles from *C. elegans* carry proteins, non-coding RNAs, miRNAs, lipid nanodomains, enzymes and other chemicals (Nikonorova *et al*., 2022; Russell *et al*, 2020). Collectively, these findings suggest the involvement of RNA in learning and memory and may allow for the hypothesis that non-coding RNAs or miRNAs could be involved in LTAM uptake from trained to naïve animals through EVs.

In summary, we report a novel mechanism of LTAM transfer through environmentally released EVs. This, to the best of our knowledge, is the first report of organic transfer of long-term associative memory between organisms and across species.

## Acknowledgements

The authors are grateful to Maureen Barr, Abhishek Bhattacharya, Björn Brembs, Sean Curran, Yogesh Dahiya, Aurnab Ghose, Patrick Laurent, Sandhya Koushika, Coleen Murphy, Aleksandra Nawrocka, Piali Sengupta, Renee Set, Jagan Srinivasan and Juan Wang for experimental suggestions or critique on the manuscript. The authors thank Deepak Nair and Shrivallabh Shankar for help with the imaging experiment and the IISc Mass Spectrometry Facility at the Division of Biological Sciences for the LC-MS data and analyses. We also thank Shrinithi Natarajan for help with the RNA sequencing experiment. We thank Maureen Barr for reagents. Some strains were provided by CGC, which is funded by NIH Office of Research Infrastructure Programs (P40 OD010440). We also thank NBRP, Japan for providing the *cil*-7 *(tm5848)* allele. All illustrations were created in BioRender. Lastly, we thank Palagiri Suresh for routine help and the members of Kavita Babu’s lab for suggestions and critique on the manuscript.

## Funding

The work was supported by an MoE STARS Grant [no. MoE/STARS-1/454] and the DBT/Welcome Trust India Alliance Fellowship [grant number IA/S/19/2/504649] awarded to KB. This work was part funded by a DBT Janaki Ammal National Women Bioscientist Award [no. BT/HRD-NBA-NWB/38/2019-20], and ANRF grants [nos. SPG/2022/000182 and CRG/2023/001950] to KB. The authors are also grateful to IISc and the DBT IISc partnership program for intramural funding. MB was supported by IISc funds, TN was supported by a Bayer MEDHA Fellowship, HN was supported by a KVPY Fellowship, and KK is supported by CSIR-JRF and CSIR-SRF fellowships. The funders had no role in experimental design, data collection or analysis, decision to publish, or preparation of the manuscript.

## Conflict of Interest

Authors declare no conflict of interests.

## Notes

### Competing Interest Statement

The authors have declared no competing interest.

### Summary of Updates

LC-MS data was verified with additional experiments.

