## Supplemental information for "Extracellular vesicles aid in the transfer of long-term associative memory between *Caenorhabditis elegans*"

#### Supplementary Information

#### Supplementary methods

##### Behavioural assay:

The learning paradigm for long-term associative memory formation in *C. elegans* was performed as previously described with minor modifications (Dahiya *et al*, 2019). The assay can be divided into two parts (training, testing), which are described below. The training paradigm is illustrated in Figure 1A.

##### Training

A dry heating block was maintained at 37°C, 20 µl of 10% isoamyl alcohol (IAA)/1% Diacetyl (diluted in ethanol) was placed on top of a glass cover slip and that was in turn placed on top of the heating block. A 60 mm petri plate containing *C. elegans* was inverted on the cover slip containing IAA/diacetyl placed on the heat block for two minutes. The training cycle was repeated 5 times with an inter-training interval of 10 minutes.

##### Testing

This phase uses chemotaxis assay to test the memory formation in all the *C. elegans* that had been trained. Trained animals were kept for 20 hours at 22°C. *C. elegans* were picked with platinum loops and allowed to crawl on unseeded Nematode Growth media (NGM) plates for about 30 seconds.

8 to 10 animals were gently transferred to the center of a 90 mm NGM agar plate (without food). 1 µl of the native chemoattractant (1% IAA/1% heptanone/0.1%diacetyl) were placed at one end of the plate as used in previous studies (Dahiya *et al.*, 2019; Zhang *et al*, 2016). Chemotaxis behaviour was observed for the next ten minutes.

To quantify the chemotaxis behaviour, the movement of *C. elegans* away or towards the chemoattractant for all the four sets of animals was considered. If the animals move up the concentration gradient of the chemoattractant, then the displacement was considered positive. However, if the animals move down the concentration gradient of the chemoattractant, then the displacement was considered negative.

**Tracking movement of *C. elegans* and its analysis:**

During the testing phase of the associative learning paradigm, the plate with 8-10 *C. elegans* carrying out chemotaxis was recorded using a ThorLabs 8051C-USB camera. The setup was illuminated from underneath, with the camera mounted above the plate, for a duration of 10 minutes each.

These videos were then analysed using FIJI, and an associated plugin, Trackmate (Ershov *et al*, 2022; Schindelin *et al*, 2012). The video files were pre-processed through FIJI's suite of contrast enhancement, thresholding and background subtraction tools. Subsequently, these background-subtracted images with only *C. elegans* visible were run through the Trackmate plugin to enable spot-detection and automated track assignment, using a time-series based clustering method called Kalman filtering (implemented by the TrackMate plugin as the Kalman Tracker method). Despite the high quality of background subtraction, due to collisions in paths or occasional noise, the tracks were manually double-checked and occasionally corrected, enabling this semi-automated pipeline to be robust to any perturbations in the images.

The output of TrackMate, track files and spot files (sorted post-tracking by the unique *C. elegans* identity of each spot, per experiment), provide a number of parameters including the entire distance covered by individual *C. elegans*, average speed of the animals in a chemotaxis assay and the x- and y-coordinates of every *C. elegans* at every frame of recording. This allows us to subsequently calculate, on a per-experiment basis, the displacement of a *C. elegans* from its final position with respect to the position of the chemoattractant and the average position of a *C. elegans* over the entire chemotaxis assay. This was achieved through the use of the Pandas library for Python with its suite of vectorized operations within its data frames, allowing the integration of the spot and track outputs and computations between them.

**Chemotaxis index:**

This is calculated by considering the distance travelled by a *C. elegans or C. briggsae* and its displacement with respect to IAA/diacetyl/heptanone at the end of the ten minutes of recording. Aversive learning leads to loss of attraction towards IAA/diacetyl/heptanone, and hence, lowers the chemotaxis index from naïve conditions. The formula is as follows:

$$\text{Chemotaxis index (CI)} = \frac{\mathit{Displacement\ along\ the\ chemoattractant\ gradient}}{\mathit{Total\ distance\ travelled}}$$

The process described above was also used with for testing the CI of *C. briggsae*.

**Supplementary Tables**:

**S1 (Strains used in this study)**

| Strain number | Genotype | Source |
|---|---|---|
| BAB9000 | *crh-1(tz2)* (CGC strain YT17) | From (Dahiya *et al.*, 2019), further outcrossed 3X |
| BAB9001 | *daf-22 (m130)* (CGC strain DR476) | Outcrossed 3X |
| PT1194 | *klp-6 (my8) III; him-5(e1490) V* | From CGC, USA |
| TM5848 | *cil-7 (tm5848) I* (shows him phenotype) | From NBRP, Japan |
| PT3602 | *cil-7 (my61[cil-7::mNG])I; him-5(e1490) V* | Gift from Maureen Barr (Wang *et al*, 2021) |
| BAB9002 | *klp-6 (my8);* P*klp-6*::GFP::KLP-6 (array, IndEx 9002) | This study |
| AF16 | *C. briggsae* G16 isolate | From CGC, USA |

**S2 (Primers used in this study)**

| Primer Id | Sequence | Primer type | mutation |
|---|---|---|---|
| YD157 | TGGAAGGAGGAGGAGATGGAAA | Genotyping (external forward) | *crh-1(tz2)* |
| YD158 | GCAGTACAGCTCTTTCAGCGTT | Genotyping (internal forward) | *crh-1(tz2)* |
| YD159 | AATTCGGCACAACGGACTGG | Genotyping (external reverse) | *crh-1(tz2)* |
| NS269 | CGGTTGCTCCGATAGGATGACT | Genotyping (SNP mutant forward) | *daf-22 (m130)* |
| NS270 | CGGTTGCTCCGATAGGATGACC | Genotyping (SNP WT forward) | *daf-22 (m130)* |
| NS262 | CCCAGTTCACCAAGGAATTCTCTC | Genotyping (SNP mutant and WT reverse) | *daf-22 (m130)* |

#### Supplementary figure legends:

**Figure S1: Track images of naïve and trained *C. elegans***

(A) Track images of WT naïve and trained *C. elegans*, each color represents an individual *C. elegans* track. The IAA spot is indicated as a dashed circle in all track images. (B) Lifecycle of *C. elegans*, showing the time point of animal transfer. (C) Violin plots indicating CI in *C. elegans* transferred at 12 hours from training after a washing step. Multiple comparisons were done using one-way ANOVA and p-values were adjusted using Dunnett's correction; "ns" indicates not significant.

**Figure S2: *C. elegans* do not gain LTAM when they are exposed to heat and IAA one after the other**

(A) Track images of WT naïve, naïve *C. elegans* added to an empty plate that had had trained *C. elegans* and trained *C. elegans* added to an empty plate that had had naïve animals (ie exchange of animals from naïve to trained plates). Each color represents an individual *C. elegans* track. (B) Schematic illustration of unpaired assays, where each cue is presented one after the other instead of simultaneously. (C, D) Violin plots indicating CI in animals trained in the unpaired assays; IAA followed by heat and heat followed by IAA, respectively. Statistical analysis for S2C was performed using Mann-Whitney's non-parametric U test. Statistical analysis for S2D was performed using one-way ANOVA and p-values were adjusted using Dunnett's correction, "ns" indicates not significant in all cases.

**Figure S3: Illustration of AWA and AWC neurons sensing different odours**

Illustration of chemosensory neuron AWC detecting IAA and heptanone and the AWA neuron detecting diacetyl.

**Figure S4: Track images of *crh-1* mutant animals**

Track images of *crh-1* naïve, *crh-1* trained, *crh-1* naïve transferred to an empty plate that had had WT trained *C. elegans* and *crh-1* trained animals transferred to an empty plate that had had WT trained *C. elegans*. Each color represents an individual *C. elegans* track.

**Figure S5: Track images of WT and *klp-6* mutant animals**

Track images of WT naïve and trained animals in the top panels, followed by track images of *klp-6* mutants naïve and trained *C. elegans* in the middle panel. The bottom panel shows track images of *klp-6* trained animals transferred to an empty plate that had had WT trained *C. elegans* and WT trained animals transferred to an empty plate that had had *klp-6* trained *C. elegans*. Each color represents an individual *C. elegans* track.

**Figure S6: Structures of molecules present exclusively on WT trained plates**

The chemical structures of eight of the nine molecules found only on the WT trained plates after LC-MS analyses are indicated here in the order that they appear in the list on Figure 7A lower panel. The images were downloaded from PubChem.

**Figure S7: Track images of *C. elegans* treated with a cocktail of chemicals obtained in through LC-MS analyses of treating animals with heat and IAA**

(A) Track images of WT naïve, WT naïve+Imazapyr, WT naïve+2M5M, WT naïve+SGCDC and WT naïve+all 3 chemicals and trained animals recorded at 3.5 hours after chemical treatment or training. The tracks indicate response of the animals to IAA. (C) WT naïve and trained animals recorded at 3.5 hours post training. (D) WT naïve and trained animals recorded at 6 hours post training.

**Figure S8: *C. briggsae* does not appear to form LTAM to IAA**

(A) Violin plots indicating CI in *C. briggsae* trained with the IAA and heat associative learning paradigm. No LTAM is seen in *C. briggsae* trained with IAA and heat. (B) Violin plots indicating CI of *C. elegans* naïve and trained, *C. briggsae* naïve, *C. briggsae* naïve transferred to an empty plate that had had trained WT *C. elegans* and *C. elegans* trained transferred to an empty plate that had had naïve *C. briggsae.* Multiple comparisons were performed using one-way ANOVA and p-values were adjusted using Dunnett's correction, "****" indicates $p< 0.0001$ and "ns" indicates not significant.

### Figure S1

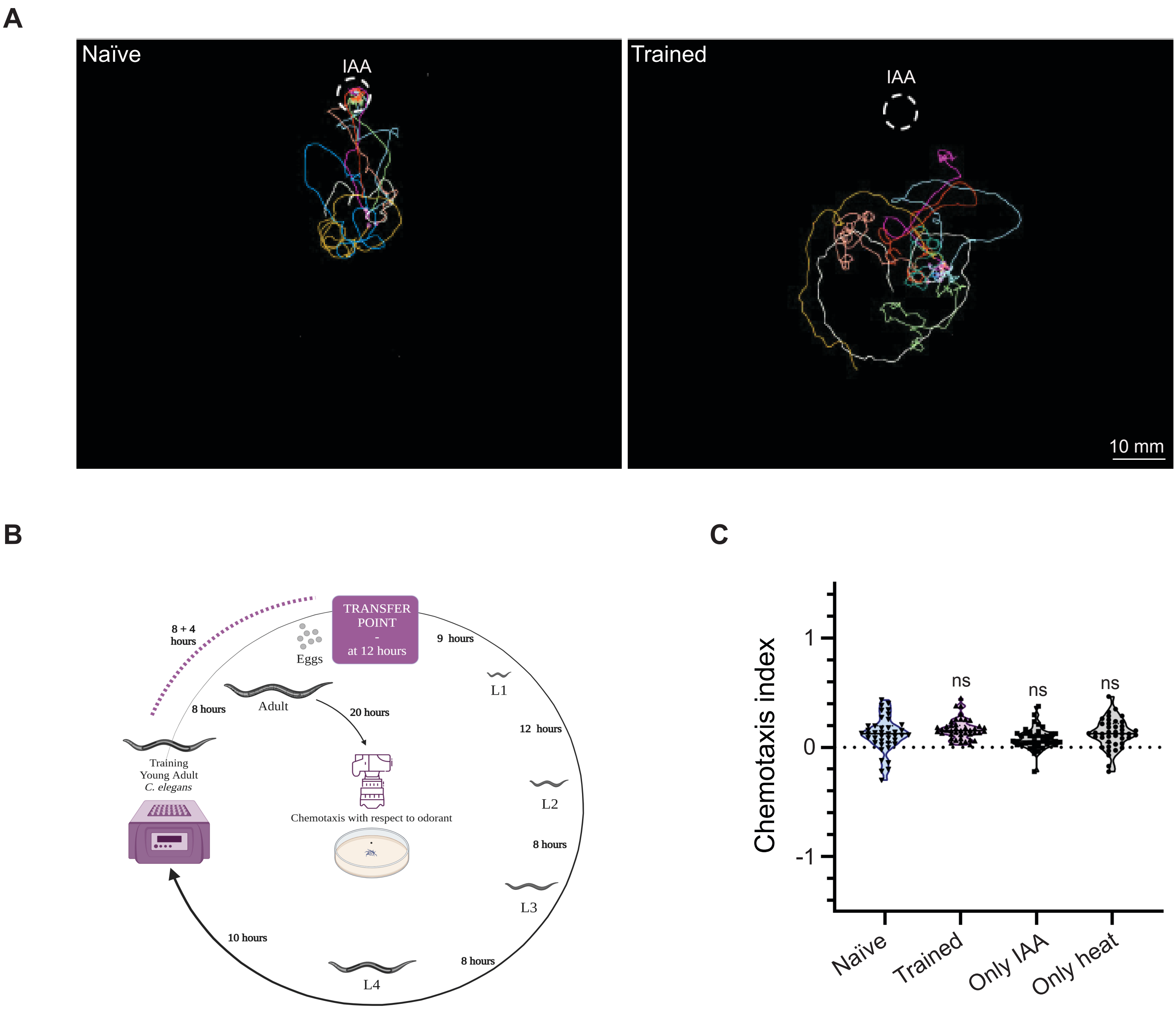
A
Naïve
IAA
Trained
IAA
10 mm
B
TRANSFER POINT
-
at 12 hours
8 + 4 hours
Eggs
9 hours
L1
8 hours
Adult
20 hours
12 hours
Training
Young Adult
C. elegans
Chemotaxis with respect to odorant
L2
8 hours
L3
10 hours
8 hours
L4
C
Chemotaxis index
1
0
-1
ns
ns
ns
Naïve
Trained
Only IAA
Only heat

#### Figure S2

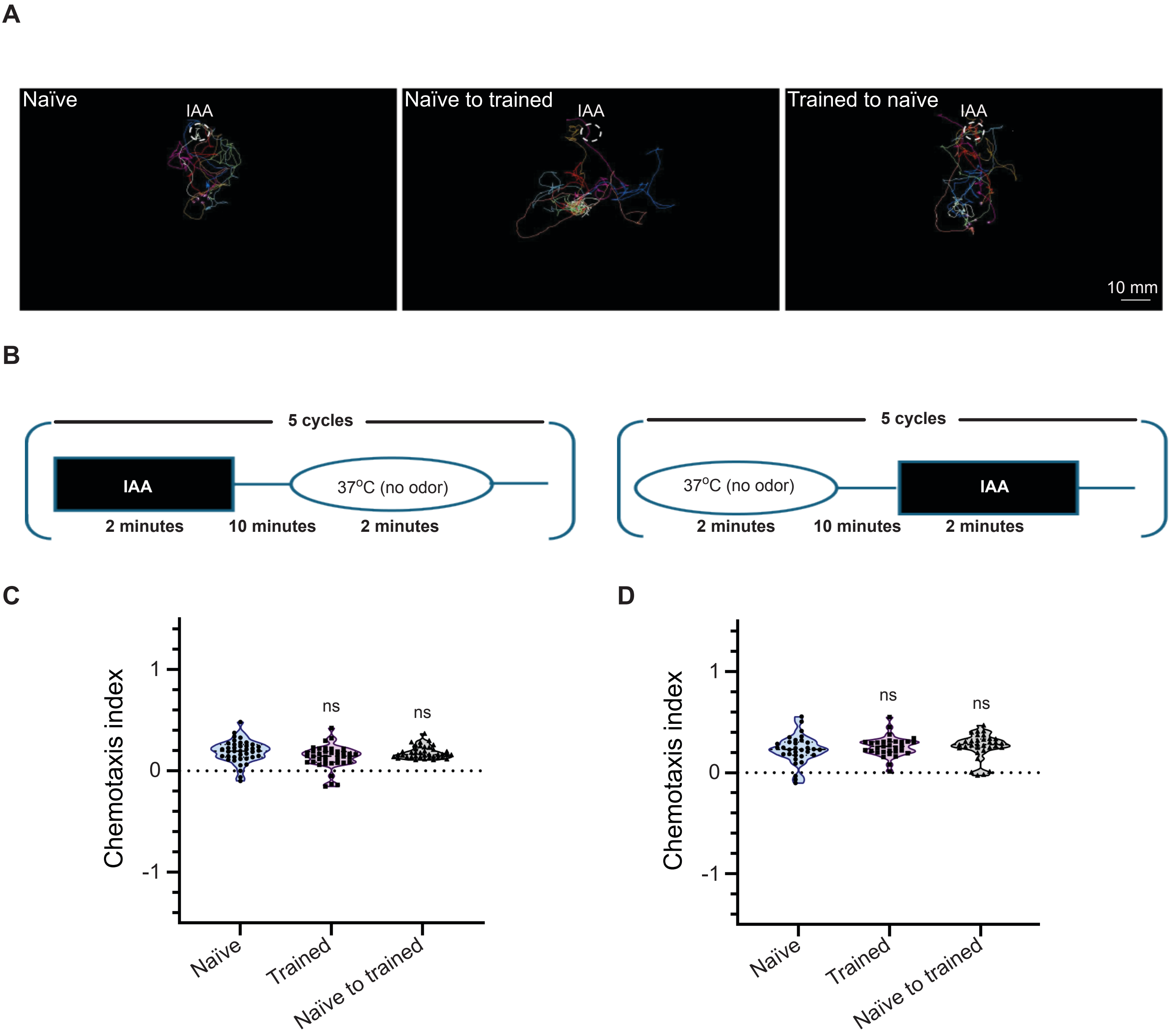
A
Naïve
IAA
Naïve to trained
IAA
Trained to naïve
IAA
10 mm
B
5 cycles
IAA
37°C (no odor)
2 minutes
10 minutes
2 minutes
5 cycles
37°C (no odor)
IAA
2 minutes
10 minutes
2 minutes
C
Chemotaxis index
1
0
-1
ns
ns
Naïve
Trained
Naïve to trained
D
Chemotaxis index
1
0
-1
ns
ns
Naïve
Trained
Naïve to trained

**Figure S3**

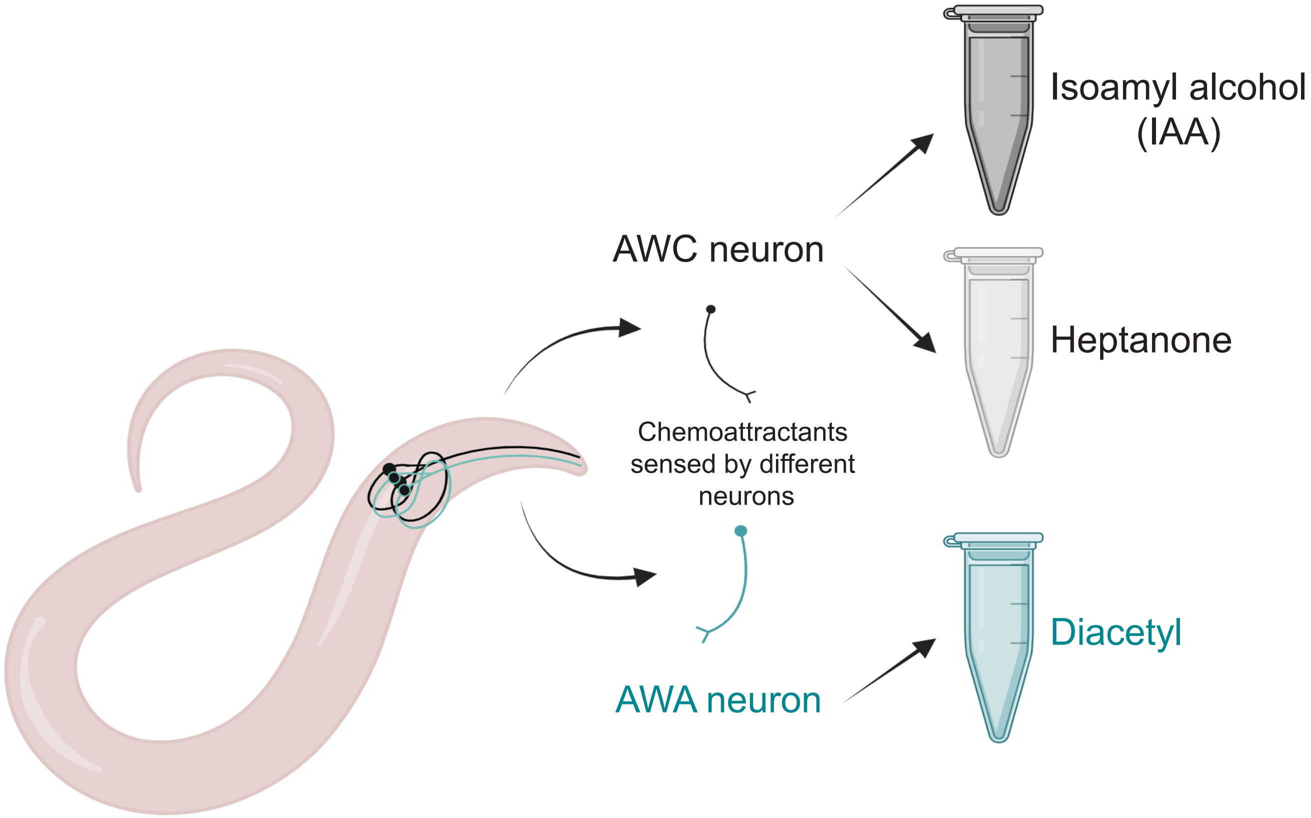
AWC neuron
Isoamyl alcohol (IAA)
Heptanone
Chemoattractants sensed by different neurons
AWA neuron
Diacetyl

### Figure S4

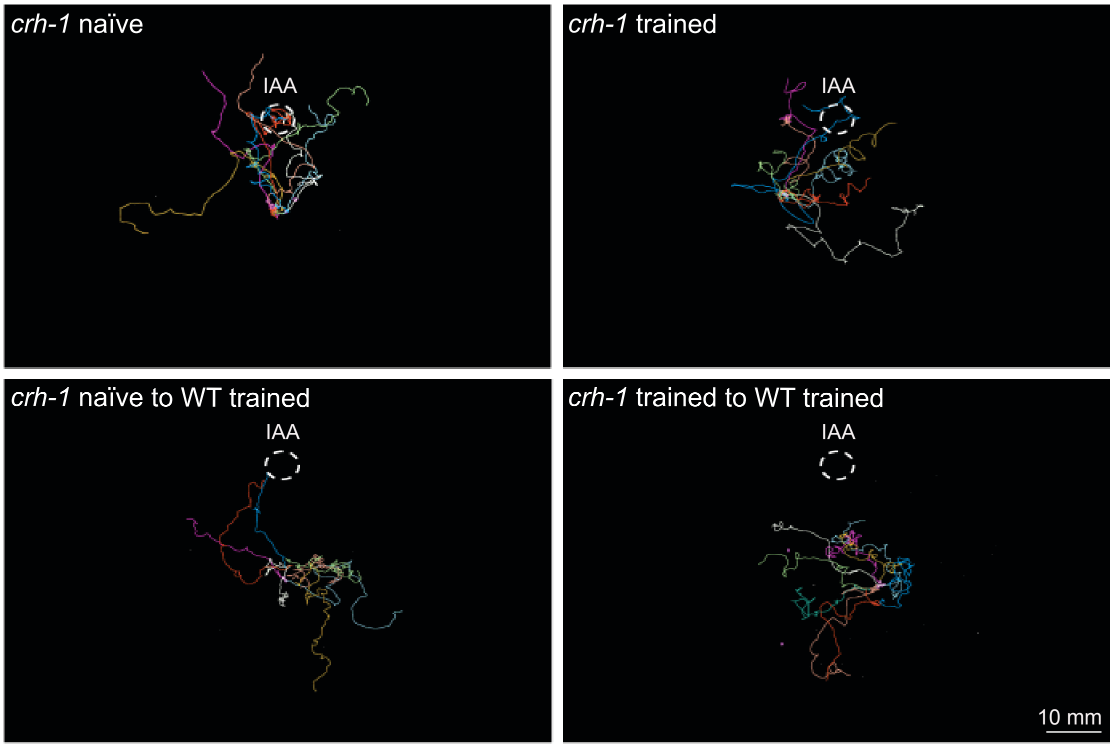

*crh-1* naïve
IAA
*crh-1* trained
IAA
*crh-1* naïve to WT trained
IAA
*crh-1* trained to WT trained
IAA
10 mm

#### Figure S5

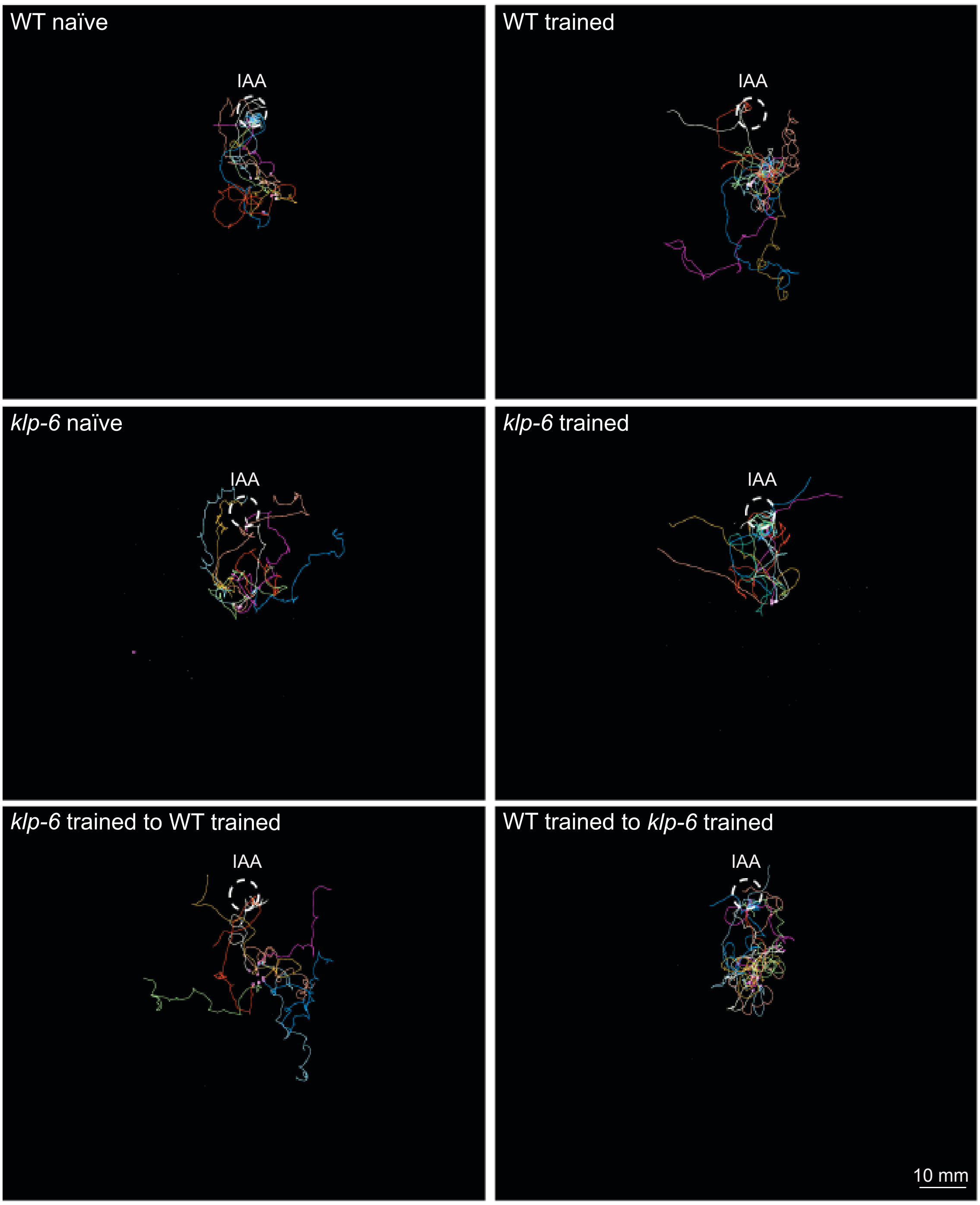

WT naïve
IAA
WT trained
IAA
*klp-6* naïve
IAA
*klp-6* trained
IAA
*klp-6* trained to WT trained
IAA
WT trained to *klp-6* trained
IAA
10 mm

### Figure S6

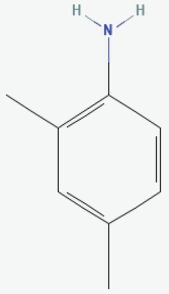

**2,4-Xylidine**

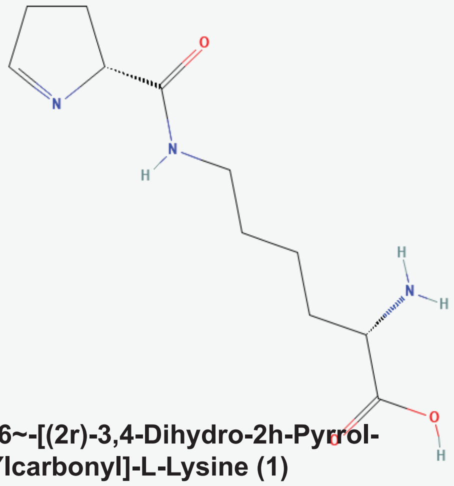

**N~6~-[(2r)-3,4-Dihydro-2h-Pyrrol-2-Ylcarbonyl]-L-Lysine (1)**

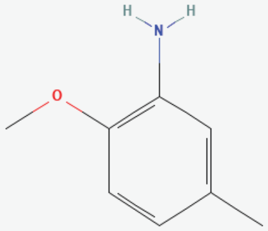

**2-Methoxy 5-methylaniline (2M5M)**

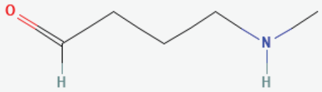

**4-(Methylamino) butanal**

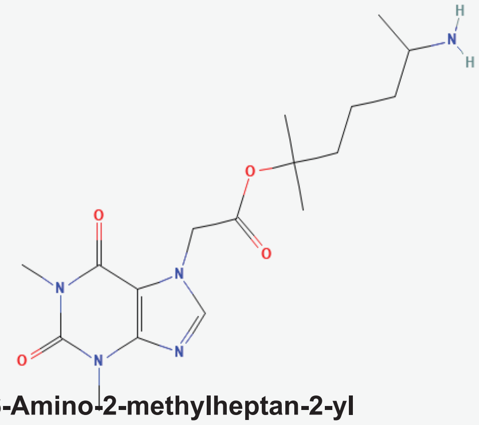

**6-Amino-2-methylheptan-2-yl (1,3-dimethyl-2,6-dioxo-1,2,3,6-tetrahydro-7H-purin-7-yl)acetate**

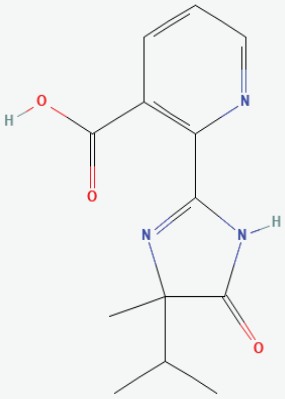

**Imazapyr**

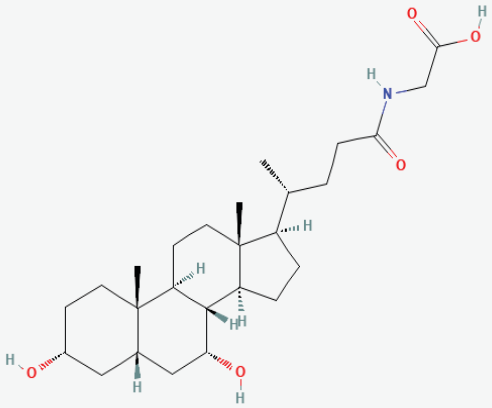

**Glycochenodeoxycholic Acid (Socium salt- SGCDC)**

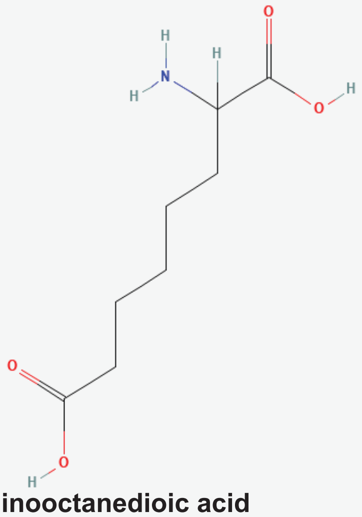

**2-Aminooctanedioic acid**

### Figure S7

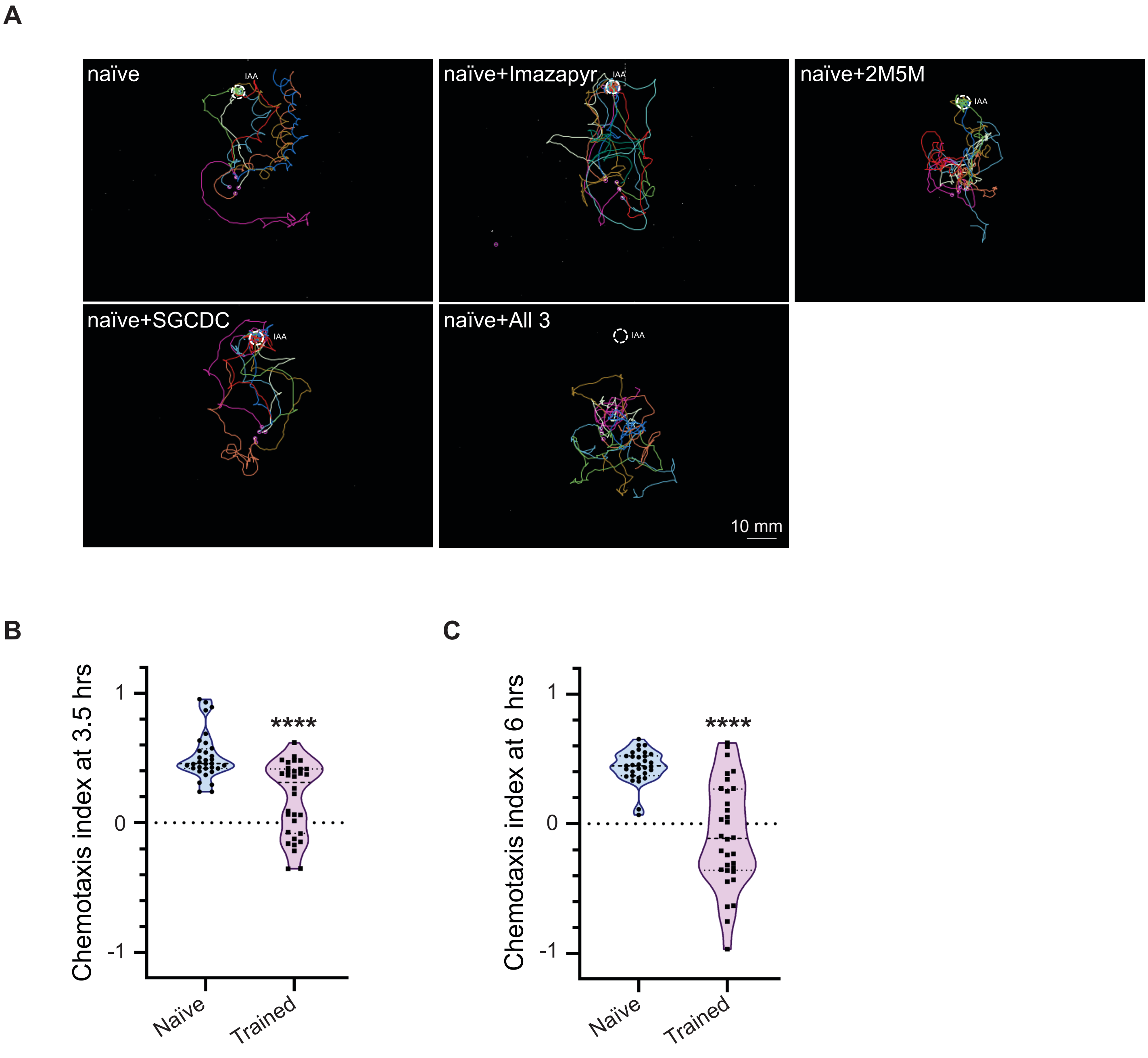
A
naïve
IAA
naïve+Imazapyr
IAA
naïve+2M5M
IAA
naïve+SGCDC
IAA
naïve+All 3
IAA
10 mm
B
Chemotaxis index at 3.5 hrs
1
0
-1
****
Naïve
Trained
C
Chemotaxis index at 6 hrs
1
0
-1
****
Naïve
Trained

#### Figure S8

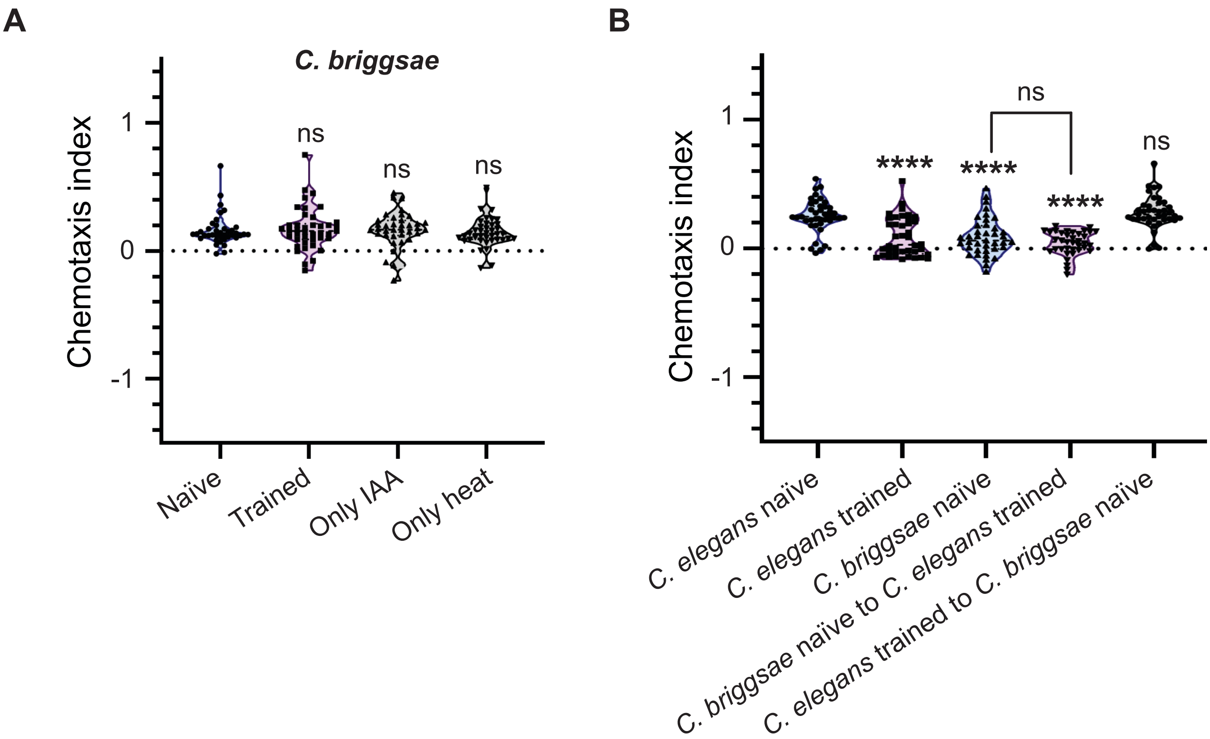
A
C. briggsae
Chemotaxis index
1
0
-1
ns
ns
ns
Naïve
Trained
Only IAA
Only heat
B
Chemotaxis index
1
0
-1
ns
****
****
****
ns
C. elegans naïve
C. elegans trained
C. briggsae naïve
C. briggsae naïve to C. elegans trained
C. elegans trained to C. briggsae naïve
